# New Targeted Approaches for Epigenetic Age Predictions

**DOI:** 10.1101/799031

**Authors:** Yang Han, Julia Franzen, Thomas Stiehl, Michael Gobs, Chao-Chung Kuo, Miloš Nikolić, Jan Hapala, Barbara Elisabeth Koop, Klaus Strathmann, Stefanie Ritz‐Timme, Wolfgang Wagner

**Author notes:** **Correspondence:** Wolfgang Wagner, M.D., Ph.D., Helmholtz-Institute for Biomedical Engineering, Stem Cell Biology and Cellular Engineering, RWTH Aachen University Medical School, Pauwelsstraße 20, 52074 Aachen, Germany.

## Abstract

Aging causes epigenetic modifications, which are utilized as a biomarker for the aging process. While genome-wide DNA methylation profiles enable robust age-predictors by integration of many age-associated CG dinucleotides (CpGs), there are various alternative approaches for targeted measurements at specific CpGs that better support standardized and cost-effective high-throughput analysis. In this study, we utilized 4,650 Illumina BeadChip datasets of blood to select the best suited CpG sites for targeted analysis. DNA methylation analysis at these sites with either pyrosequencing or droplet digital PCR (ddPCR) revealed a high correlation with chronological age. In comparison, bisulfite barcoded amplicon sequencing (BBA-seq) gave slightly lower precision at individual CpGs. However, BBA-seq data revealed that the correlation of methylation levels with age at neighboring CpG sites follows a bell-shaped curve, often accompanied by a CTCF binding site at the peak. We demonstrate that within individual BBA-seq reads the DNA methylation at neighboring CpGs is not coherently modified but reveals a stochastic pattern. Based on this, we have developed an alternative model for epigenetic age predictions based on the binary sequel of methylated and non-methylated sites in individual reads, which reflects heterogeneity in epigenetic aging within a sample. Thus, the stochastic evolution of age-associated DNA methylation patterns, which seems to resemble epigenetic drift, enables epigenetic clocks for individual DNA strands.

## Introduction

During aging, DNA methylation (DNAm) is continuously lost or gained at specific CG nucleotides (CpG sites) of our genome. Conversely, the DNAm levels at multiple CpG sites can be combined to estimate age, and these models are often referred to as “epigenetic clocks” (Field et al. 2018). Epigenetic clocks raise hopes as a biomarker in forensic medicine, to determine donor age of an unknown specimen or of a person with allegedly unknown age (Horvath and Raj 2018). On the other hand, accelerated epigenetic aging has been shown to be associated with shorter life expectancy (Marioni et al. 2015; Lin et al. 2016; Zhang et al. 2017; Lu et al. 2019; Lund et al. 2019), and it is liable to be affected by environmental exposure, gender, specific mutations and diseases (Fiorito et al. 2019; Jeffries et al. 2019; Martin-Herranz et al. 2019). Therefore, epigenetic clocks seem to reflect aspects of biological age, which opens perspectives as a surrogate for intervention studies. It is even conceivable that therapeutic regimen in future medicine will rather be stratified by epigenetic age than chronological age. To translate epigenetic biomarkers into an approved medical test, it is advantageous to select a manageable set of informative genomic regions, which can be targeted by DNA methylation assays that are sufficiently fast, cheap, robust and widely available for clinical diagnostics (Kristensen and Hansen 2009; Blueprint-consortium 2016).

Initially, models for epigenetic age predictions were based on Illumina BeadChip data (Bocklandt et al. 2011; Koch and Wagner 2011). This microarray platform enables cost-effective and relatively precise DNAm measurements at single-base resolution. In contrast, whole genome bisulfite sequencing (WGBS) or reduced representation bisulfite sequencing (RRBS), does not always cover the same CpG sites and a limited number of reads may entail lower precision of DNAm levels (Walker et al. 2015; Wagner 2017). Another advantage of the Illumina BeadChip technology is that a multitude of publicly available datasets can easily be integrated into the analysis. Various different epigenetic age-predictors have been described that consider up to several hundreds of CpGs (Hannum et al. 2013; Horvath 2013; Weidner et al. 2014). The most commonly used age predictor for multiple different tissues is based on 353 CpGs to facilitate epigenetic age predictions with a median error of 3.6 years (Horvath 2013). However, this predictor was trained for the 27k and 450k platform and is not applicable to the current version of the EPIC Illumina BeadChip array.

For targeted analysis of specific age-associated CpGs, various studies described epigenetic age-predictors based on bisulfite pyrosequencing (Weidner et al. 2014; Zbieć-Piekarska et al. 2015) or by the Sequenom’s EpiTYPER assay (Garagnani et al. 2012). Single Base Primer Extension Assay (SNaPshot) was also used by many laboratories for epigenetic age prediction, but the accuracy is apparently lower (Lee et al. 2015; Hong et al. 2017). Recently, droplet digital PCR (ddPCR) was reported to enable precise DNAm measurements (Yu et al. 2015; Zemmour et al. 2018), and hence it might facilitate epigenetic age predictions without PCR bias. Barcoded bisulfite amplicon sequencing (BBA-seq), which is based on next generation sequencing, enables multiplexed analysis of PCR amplicons (Bernstein et al. 2015; Theophilou et al. 2015). The strength of BBA-seq lies within the parallelization of multiple DNA sequences on one lane with relatively long amplicons (up to 500 bases), and with very high coverage (usually > 1000 fold) (Smith et al. 2010; Franzen et al. 2017). So far, only few groups described ddPCR (Shi et al. 2018) and BBA-seq (Naue et al. 2017; Aliferi et al. 2018) for epigenetic age predictions and a direct comparison of these methods is still elusive.

It is largely unclear, how age-associated DNAm is regulated and if it is functionally relevant, *per se*. Transcription factors (TFs) or long non-coding RNAs (lncRNAs) might target epigenetic writers, such as DNA methyltransferases (DNMTs) or ten-eleven translocation family enzymes (TETs), to specific sites in the genome (Kalwa et al. 2016), This process may also involve alternative splicing of DNMTs (Bozic et al. 2018). CCCTC-binding factor (CTCF), as an insulator TFs that is involved in chromatin architecture, has also been shown to be methylation sensitive (Wiehle et al. 2019). If age-associated DNAm was directly regulated by epigenetic writers, it would be anticipated that the DNAm pattern of neighboring CpGs on the same DNA strand is coherently modified. Alternatively, it has been proposed that age-associated DNAm is evoked by “epigenetic drift”, which may occur due to stochastic accumulation of errors, e.g. in copying DNAm patterns during cell replication (Fraga et al. 2005; Issa 2014; Zampieri et al. 2015). In this case, DNAm on individual DNA strands might rather follow stochastic patterns.

In this study, we have further optimized and compared epigenetic age predictors based on pyrosequencing, ddPCR and BBA-seq of specific age-associated regions. Furthermore, our data indicate that the correlation of age-associated DNAm with chronological age peaks at CTCF binding sites. Age-associated DNAm is not coherently modified on individual DNA strands and this enabled alternative single-read age-predictors that reveal heterogeneity in epigenetic aging within a specimen.

## Results

### Selection of age-associated CpGs for blood

To select age-associated CpGs for further comparison, we used a training set of 976 DNAm profiles of healthy human blood samples (1 to 101 years old), which are derived from seven different studies and based on the 450k Illumina BeadChip platform (Supplemental Figure S1A). To reduce the impact of leukocyte composition (Jaffe and Irizarry 2014) we excluded CpGs with high variation in DNAm across hematopoietic subsets (R > 0.02 in six different cellular subsets; Supplemental Figure S1B) (Reinius et al. 2012). Furthermore, we excluded CpGs on sex chromosomes, CpGs that are significantly affected by smoking (Gao et al. 2015; Teschendorff et al. 2015), and those which were no more comprised on the new EPIC Illumina BeadChip microarray. Thus, 416,807 CpGs were used for further selection of age-associated CpGs (Supplemental Figure S1C). To identify candidates that are best suited for targeted DNAm analysis, we selected CpGs by linear correlation with chronological age using Pearson correlation > 0.5 or < −0.5. Age-associated DNAm might also follow a non-linear monotonic function and therefore we alternatively used Spearman’s rank correlation > 0.5 or < −0.5. Since age-associated DNAm was shown to rather follow a logarithmic pattern, particularly in children (Alisch et al. 2012; Horvath 2013), we also selected CpGs that correlated with logarithmic age across all samples (Pearson correlation > 0.5 or < −0.5). Overall, 66 CpGs passed at least one of these three filter criteria – and despite the different filter criteria the overlap was remarkably high – with 19 CpGs passing all three thresholds (Figure 1A). Subsequently, we trained a multivariable model based on the 66 CpGs using functions of transformed chronological age with logarithmic dependence for pediatric and linear dependence for adult donors, as previously described by Horvath et. al (Horvath 2013). This model provided a high correlation with chronological age in the training set (R^2^ = 0.95; median error = 3.0 years; Figure 1B) and in an independent validation set of 3,674 blood samples of five different studies (R^2^ = 0.82; median error = 3.3 years; Figure 1C).

**Figure 1.**
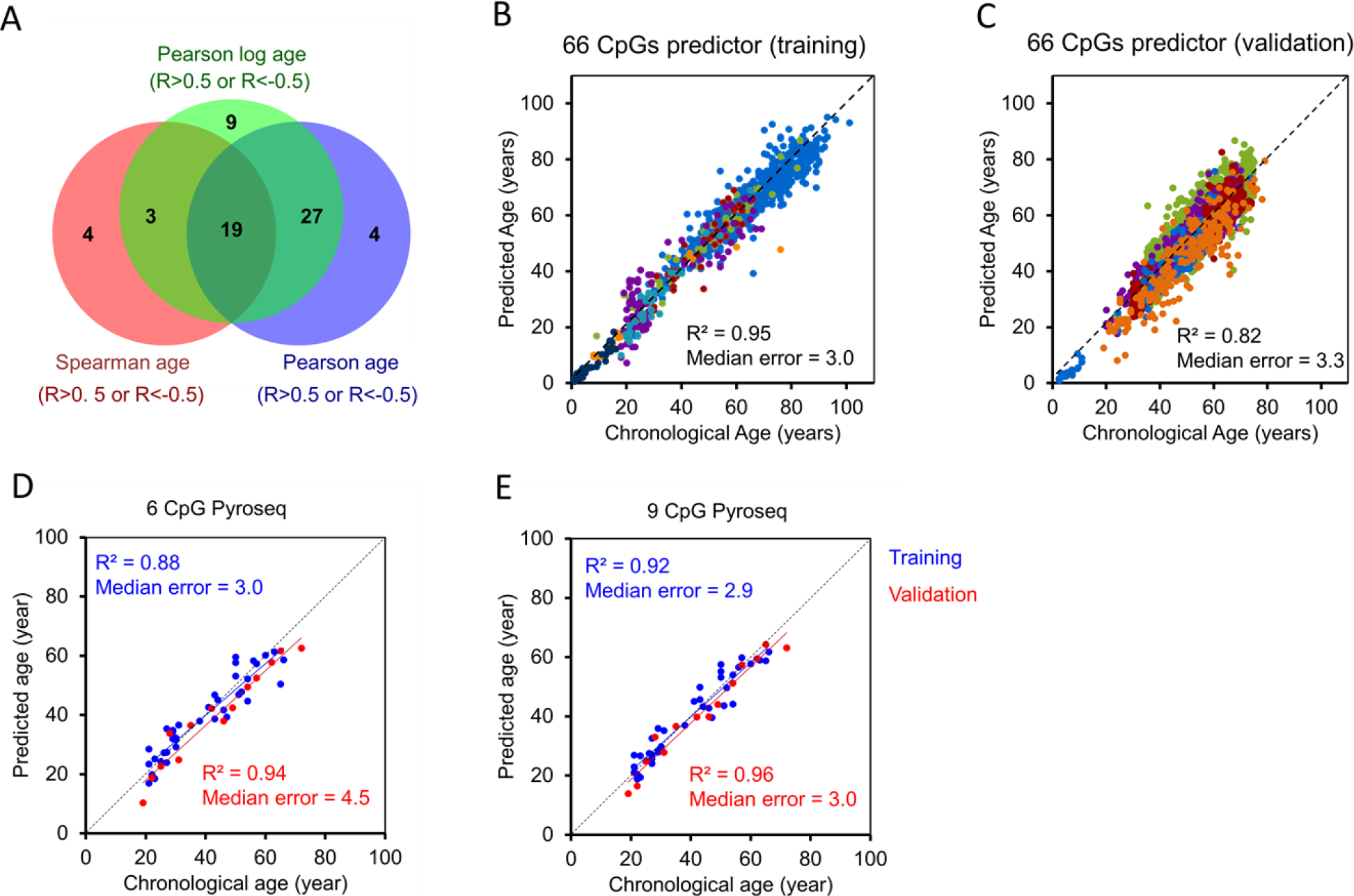
Selection of age-associated CpGs and targeted analysis with pyrosequencing. **(A)** Illumina BeadChip profiles of 976 blood samples of 7 studies (all 450k) were used to select age-associated CpGs by Pearson correlation (blue), Spearman’s rank correlation (red), or Pearson correlation after logarithmic transformation of age (green). 66 CpGs passed at least one of these thresholds (Correlation Coefficient: R>0.5 or R<−0.5) and the Venn diagram depicts a very high overlap. **(B, C)** A multivariable linear model for these 66 age-related CpGs revealed high correlation with chronological age in the training set (B; n = 976), and in an independent validation set (C; n = 3,674; color code corresponds to different studies as indicated in Supplemental Figure S1A). **(D, E)** Epigenetic age prediction based on pyrosequencing of 6 CpGs (D) and 9 CpGs (E) in blood samples of the training set (n = 40; blue) and an independent validation set (n = 14; red).

### Epigenetic age predictor based on pyrosequencing of specific CpGs

To further narrow down the best suited CpGs for targeted DNAm measurement, we selected amongst the 66 age-related CpGs those with the best linear correlation and highest slope in Illumina BeadChip training sets (Supplemental Figure S2A). The selected CpGs were associated with the genes of Elongation Of Very Long Chain Fatty Acids Protein 2 (*ELOVL2*), which is well known to reveal high age-association (Garagnani et al. 2012), Coiled-Coil Domain-Containing Protein 102B (*CCDC102B*), Four And A Half LIM Domains Protein 2 (*FHL2*), Immunoglobulin Superfamily Member 11 (*IGSF11*), Collagen Type I Alpha 1 Chain (*COL1A1*), and MEIS1 Antisense RNA 3 (*MEIS1-AS3*). When we analyzed DNAm at these CpGs in 40 blood samples by pyrosequencing, we observed overall a high correlation with chronological age, which was significantly higher than for the independent validation set of 450k Illumina BeadChip profiles (Supplemental Figure S2B). A multivariable linear regression model based on pyrosequencing of 40 samples revealed a good correlation with chronological age in an independent validation set of 14 blood samples (R^2^ = 0.94; median error = 4.5 years; Figure 1D). The accuracy was even increased, when we integrated three additional CpGs, which were selected in our previous work (associated with the genes *ITGA2B*, *ASPA* and *PDE4C* (Weidner et al. 2014)), into the model (R^2^ = 0.96; median error = 3.0 years; Figure 1E).

### Analysis of age-associated DNAm with droplet digital PCR

Droplet digital PCR (ddPCR) is based on randomly distributing a single sample into small droplets, which are then processed for PCR amplification separately. We anticipated that this technology reduces PCR bias, because sequences are amplified with different efficiency and therefore affect the ratio in conventional PCR. For each amplicon two probes were designed that either detect the methylated or the non-methylated sequence and DNAm level was subsequently determined by Poisson distribution (Figure 2A). Reliable ddPCR assays could be established for the sequences of *CCDC102B*, *COL1A1*, *MEIS1-AS3*, *FHL2, PDE4C, ASPA and IGSF11,* while this was hampered for *ITGA2B* and *ELOVL2* due to neighboring CpG sites. The ddPCR results revealed a clear correlation with chronological age at all tested CpGs, which was often even slightly higher than for pyrosequencing (Figure 2B-H). We then generated a multivariable model based on ddPCR measurement of these seven CpGs. This provided relatively reliable age-predictions in an independent validation set of 27 blood samples (R^2^ = 0.79; median error = 3.4 years; Figure 2I).

**Figure 2.**
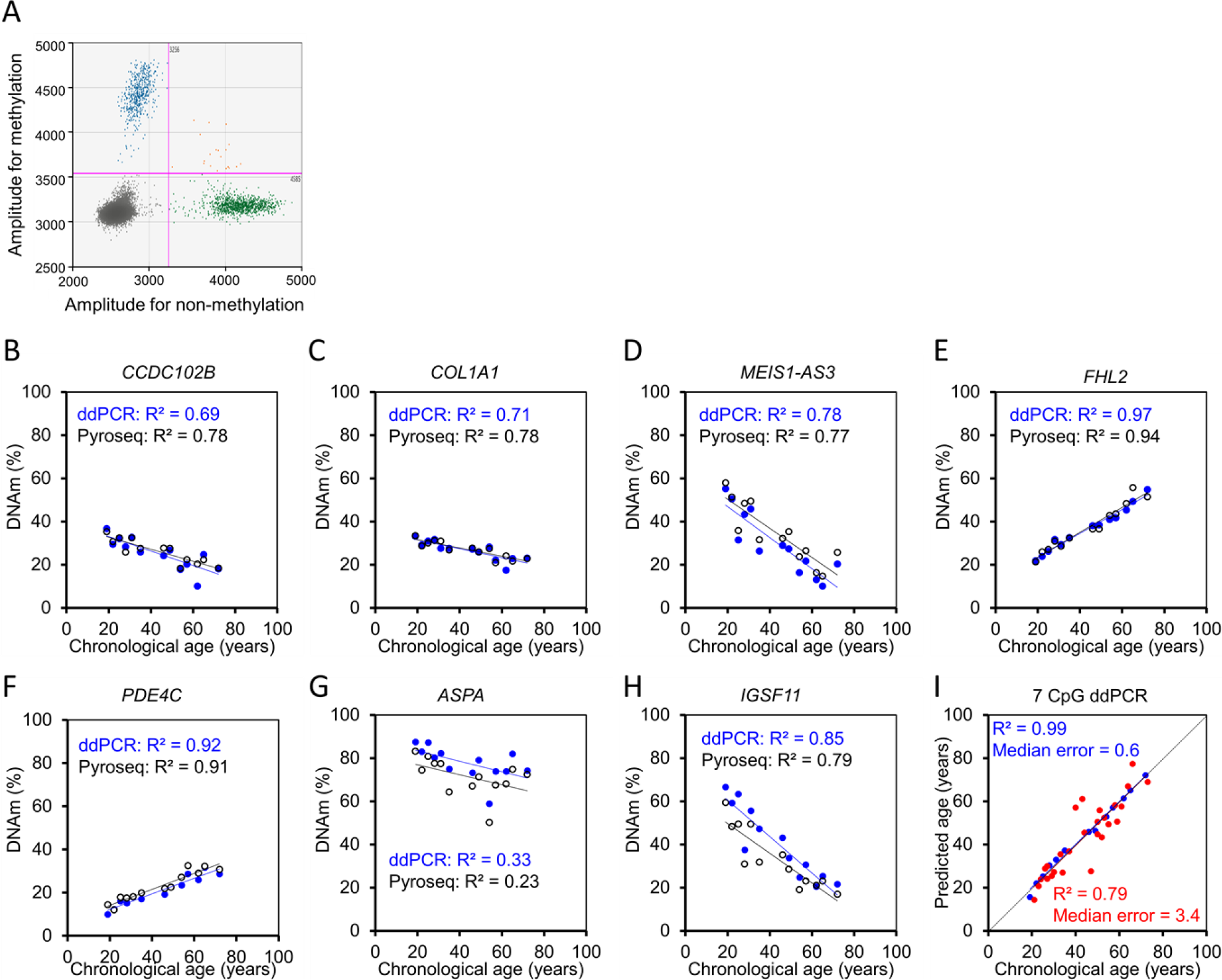
Age-associated DNA methylation measurements with droplet digital PCR. **(A)** Two-dimensional amplitude analysis of duplex ddPCR (blue: positive droplets for methylated CCDC102B; green: positive droplets for non-methylated CCDC102B; orange: double-positive droplets; black: negative droplets). **(B-H)** DNAm measurements by ddPCR versus pyrosequencing were compared for 13 blood samples. **(I**) Epigenetic age prediction based on ddPCR measurements of 7 CpGs in blood samples of the training set (n = 13; blue) and an independent validation set (n = 27; red).

### Epigenetic age-predictions with bisulfite barcoded amplicon sequencing

Subsequently, we analyzed DNAm of the nine age-associated regions with deep sequencing technology (Illumina MiSeq). Bisulfite barcoded amplicon sequencing (BBA-seq) was initially performed on a training set of 38 blood samples. Overall, DNAm levels of BBA-seq and pyrosequencing revealed good correlation (R^2^ = 0.92; Supplemental Figure S3), indicating that DNAm measurements were feasible for all amplicons. However, the correlation of DNAm levels in BBA-seq with chronological age was slightly lower as compared to pyrosequencing or ddPCR (Supplemental Figure S4), indicating that the method is slightly less precise on single CpG level. A multivariable linear regression model was then established for the nine CpGs with the highest correlation with chronological age per amplicon. This approach provided high accuracy for age-predictions in the training set (R^2^ = 0.96; median error = 3.0 years) and in an independent validation set of 39 blood samples (R^2^ = 0.86; median error = 1.9 years; Figure 3A). Alternatively, we used all CpGs within the nine amplicons for Lasso (Figure 3B) and elastic net regression models (Figure 3C), with 10-fold cross validation to prevent data overfitting. Both machine learning approaches performed better than the linear regression model on the training set, but the precision was lower on the validation set, which might be attributed to small technical offsets between different BBA-seq runs.

**Figure 3.**
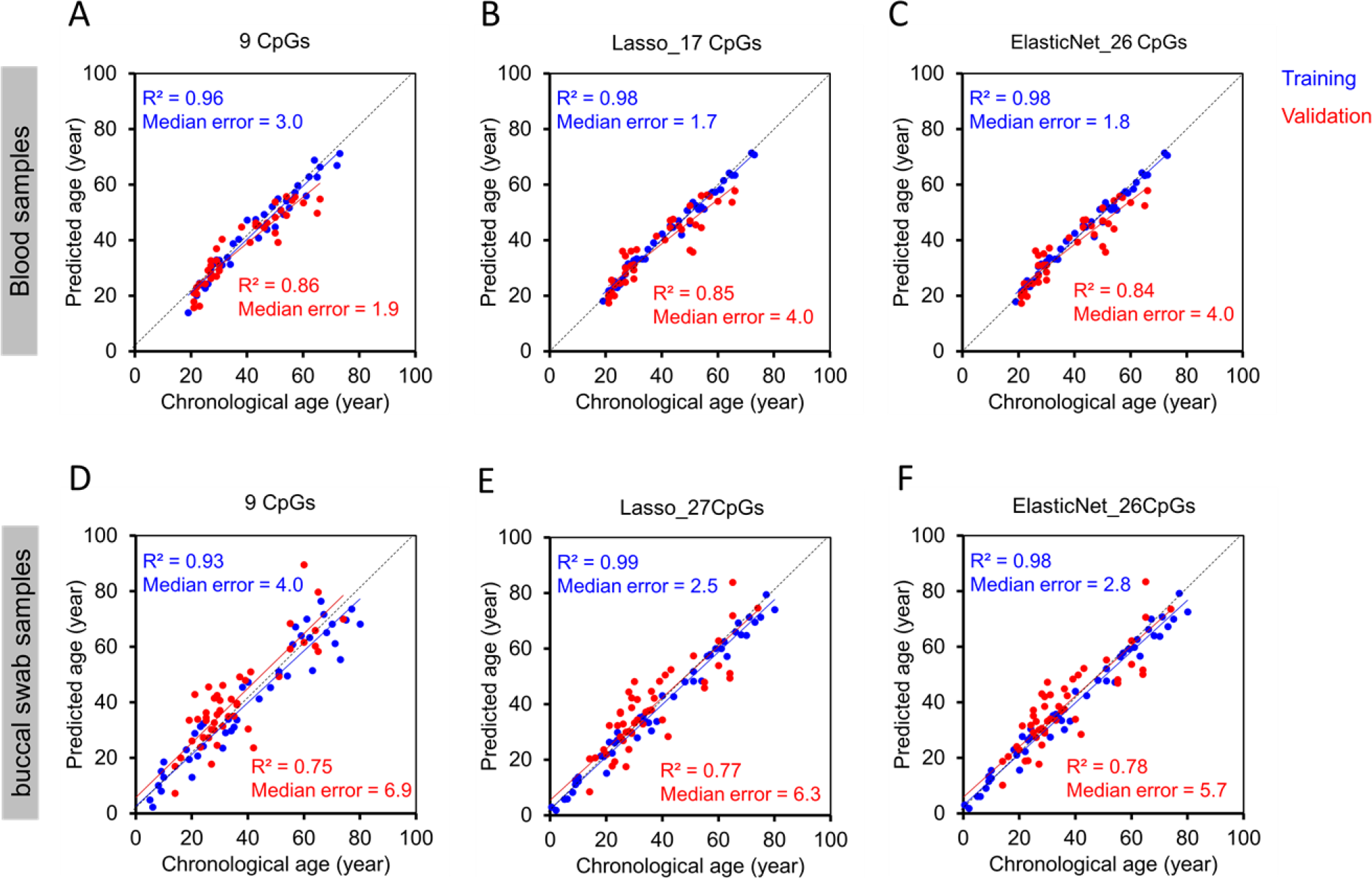
Epigenetic age predictions using BBA-seq. **(A-C)** Nine amplicons with age-associated CpGs were analyzed by bisulfite barcoded amplicon sequencing (BBA-seq) in a training set of 38 blood samples (blue) and an independent validation set of 39 samples (red). Age predictions were based on a multivariable linear model of 9 CpGs within 9 amplicons (A), Lasso regression model of 17 CpGs within 8 amplicons (B), or elastic net regression model of 26 CpGs within 8 amplicons (C). **(D-E)** In analogy, genomic DNA was isolated from buccal swabs and analyzed by BBA-seq (training set: n = 46, blue; independent validation set: n = 49, red) with a multivariable model of 9 CpGs (D), by Lasso regression model of 27 CpGs within 7 amplicons (E), and by elastic net regression model of 26 CpGs within 7 amplicons (F).

We have then analyzed, if the method is also applicable for buccal swab samples, which are a widely used specimen in legal medicine. Due to the different cellular composition, the multivariable model for the 9 CpGs was retrained to provide a good correlation in a training set of 46 buccal swab samples (R^2^ = 0.93; median error = 4.0 years), albeit the correlation remained lower in the independent validation set of 49 buccal swab samples (R^2^ = 0.75; median error = 6.9 years; Figure 3D). The accuracy could be slightly increased by Lasso (Figure 3E) and elastic net algorithms (Figure 3F) that were generated based on all CpGs of the amplicons, but it remained lower than for blood samples. This might be due to the heterogeneous composition of buccal epithelial cells and leukocytes in buccal swab samples (Eipel et al. 2016).

### Age-associated DNA methylation changes peak at CTCF binding sites

One of the advantages of BBA-seq is that amplicons are longer than in pyrosequencing and include measurement of more neighboring CpGs. We have compared the correlation of neighboring CpGs with chronological age, particularly in the amplicons of *ELOVL2*, *PDE4C* and *FHL2*, which harbor 36, 26, and 18 CpGs, respectively. Plotting of correlation coefficients against the genomic locations revealed curvy distributions (Figure 4A). Similar distributions, but less distinct, were also observed for the BBA-seq data of buccal swabs (Supplemental Figure S5). Notably, age-associated DNAm changes peaked at binding sites of CCCTC-binding factor (CTCF), which is involved in organization of chromatin structure. Furthermore, chromatin immune precipitation (ChIP)-seq data of human embryonic stem cells (hESC; GSM822297), K562 (GSM822311) and A549 cell lines (GSM822289) indicated that CTCF binds particularly at the peak of age-associated DNAm changes (Figure 4B). When we analyzed enrichment of our previously identified 66 age-associated CpGs within ChIP-Seq read peaks of CTCF, they were significantly enriched in each of the three ChIP-seq experiments (Figure 4C). These data support the notion that regulation of age-associated DNAm changes is related to CTCF binding and/or the three-dimensional chromatin conformation (Ong and Corces 2014).

**Figure 4.**
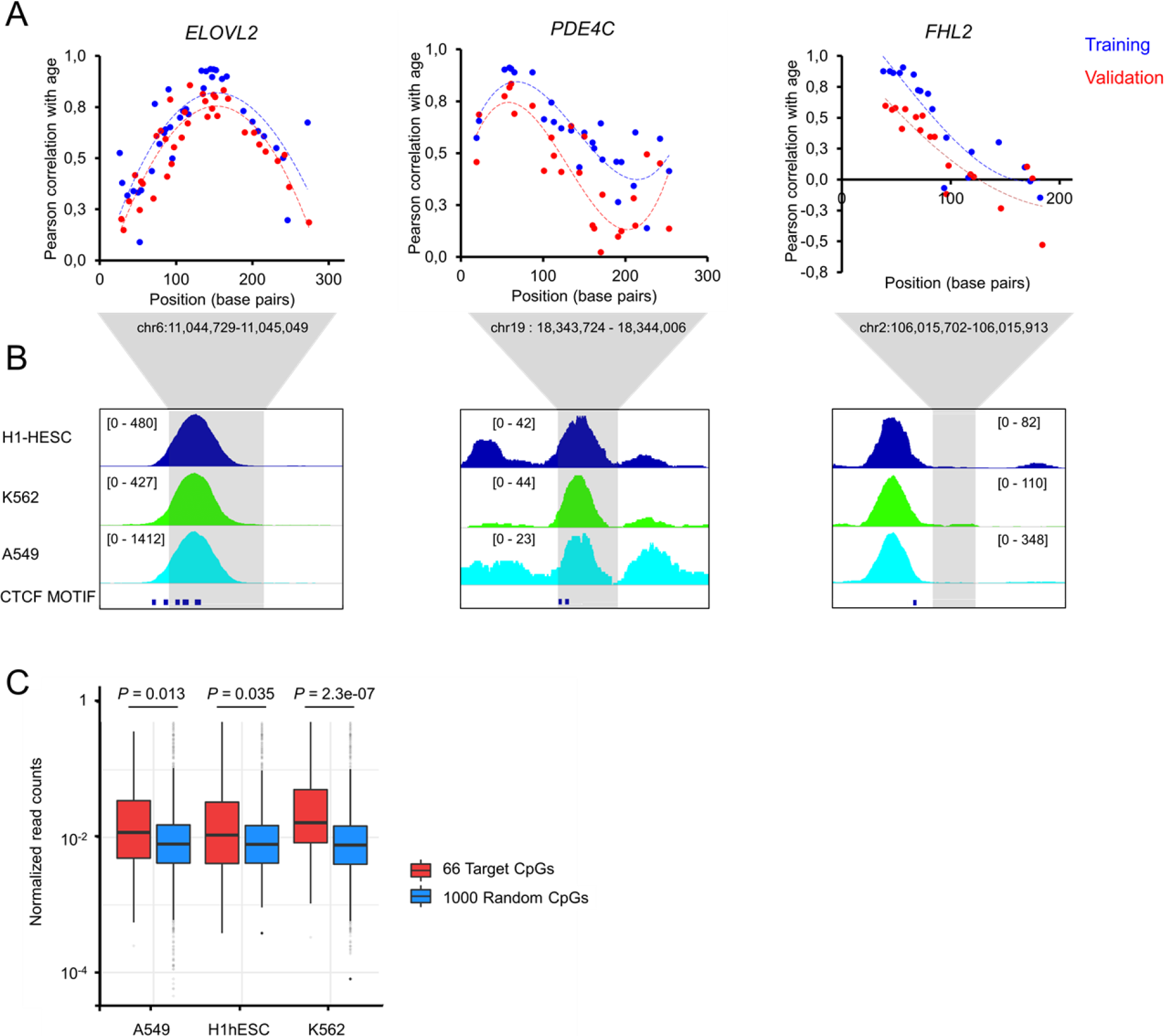
Age-associated DNAm changes peak at CTCF binding sites. **(A)** Pearson correlation of age with DNAm levels of CpGs within the amplicons of ELOVL2, PDE4C and FHL2 are plotted for the blood samples of the training set (n = 38, blue) and validation set (n = 39, red). X-axis represents the position of CpGs within the amplicons. **(B)** Enrichment of CTCF binding at the position of these amplicons (grey shaded region) was then analyzed in chromatin immune precipitation (ChIP) sequencing data of hESC (GSM822297), K562 (GSM822311) and A549 cells (GSM822289). Peak heights were automatically trimmed by IGV tool (indicated in brackets). The positions of predicted CTCF binding motives are also presented. **(C)** Boxplot of normalized read counts of CTCF ChIP-seq data of A549, hESC and K562 cell lines either at the 66 age-associated CpGs or at 1000 randomly chosen CpGs from 450k BeadChip array. Read counts from ChIP-seq data were normalized by quantile normalization and analyzed within a window of 500 base pairs (p-value was estimated by Mann-Whitney rank test).

### Analysis of age-associated DNAm patterns within individual BBA-seq reads

Age-associated DNAm peaks at specific sites in the genome, but it remained unclear whether neighboring CpGs are coherently modified or not. If DNAm was regulated by a targeted protein complex, it would be expected that neighboring CpGs are conjointly modified on individual strands. To address this question, we analyzed the DNAm pattern of individual reads in BBA-seq, which reflect the binary code of methylated and non-methylated cytosines in individual DNA strands. In fact, the DNAm patterns at the age-associated regions were very heterogeneous (Figure 5A). When we simulated random DNAm patterns based on the likeliness of DNAm at specific CpGs at a given age, the patterns were similar to the experimental results (Figure 5B). The correlation in DNAm at individual CpGs was overall very low (Figure 5C). Thus, age-associated DNAm at neighboring CpGs is apparently not coherently modified and rather seems to be acquired in a stochastic manner.

**Figure 5.**
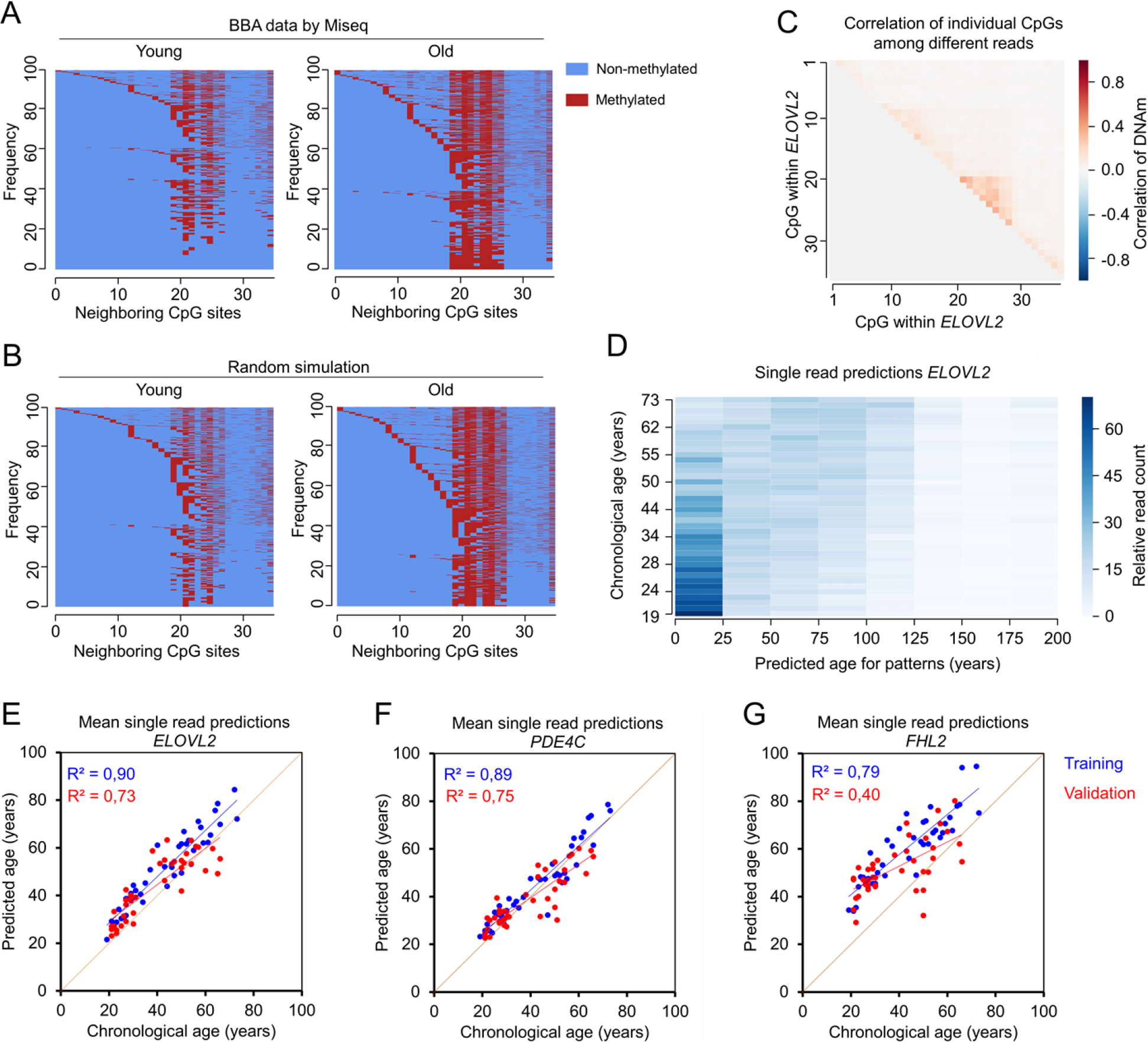
Analysis of age-associated DNAm patterns within individual BBA-seq reads. **(A)** Heat map to exemplarily depict frequencies of DNAm patterns within the 36 neighboring CpGs of the ELOVL2 amplicon in BBA-seq data of a young (21 years old) and old sample (72 years old). **(B)** For comparison, heatmaps are presented based on random simulation of DNAm patterns under the assumption that DNAm at neighboring CpGs occurs entirely independent (simulations correspond to 21 and 72 year old donors). **(C)** Pearson correlation of DNAm levels between neighboring CpG sites within ELOVL2 amplicon (BBA-seq data of training set). **(D)** For each BBA-seq read of ELOVL2 training set, we estimated the epigenetic age based on the binary sequel of methylated and non-methylated CpGs. The plot depicts relative read count of every donor in the training set that were classified for predicted ages between 0 and 200 years (Relative read count normalized by read count per sample). **(E-G)** The mean age-predictions based on individual BBA-seq reads were determined for each sample and then plotted against the chronological age of the samples of the training (blue, n = 38) and validation set (red, n = 39). This analysis was performed independently for the amplicons of ELOVL2 (E), PDE4C (F) and FHL2 (G).

We then reasoned, that DNAm patterns within individual reads might also be used for age-predictions. To this end, we developed a mathematical model based on the BBA-seq data by assigning each DNAm pattern the most likely corresponding age (between zero and 200 years; Figure 5D). As anticipated for younger donors, the model revealed a higher number of young read predictions, whereas older donors had more reads that were predicted older. Notably, the mean of strand-specific age-predictions correlated well with the chronological age of the donors in the training and validation sets (Figure 5E-G). This supports the notion that epigenetic clocks tick independently within cells of the same sample.

## Discussion

In the advent of new technologies for DNAm measurements, there is a continuous need to revisit, optimize, and validate epigenetic clocks. In this study, we used the linear correlation of age-associated DNAm with chronological age as a proxy for the precision of DNAm measurements. While all methods tested revealed good or even very good correlations with chronological age, they all have their advantages and limitations for the application in epigenetic clocks.

So far, most epigenetic clocks for human samples were derived from Illumina BeadChip datasets (Wagner 2017). It provides unparalleled opportunity to measure DNAm at single-base resolution, albeit not all CpGs of the genome are represented on these platforms (for instance about 1.7% of the human CpGs are covered by the 450k BeadChip). Our analysis clearly demonstrated that DNAm levels in 450k Illumina BeadChip data are highly correlated with chronological age at various specific CpGs. For development of our targeted assays, we focused particularly on blood samples and filtered for CpGs with low variation between leukocyte subsets to reduce the impact of the cellular composition. Our predictor with 66 age-associated CpGs provides similar, or even better accuracy than other commonly used predictors (Hannum et al. 2013; Horvath 2013; Weidner et al. 2014). Furthermore, our signature is also applicable for the current EPIC BeadChip version. While normalization regimen, integration of more CpGs, and machine learning algorithms may further enhance age-predictions, the goal of our selection was to identify suitable CpGs for targeted approaches. Focusing on a smaller number of CpGs in targeted assays is a tradeoff between the applicability and accuracy (Wagner 2017). Targeted analysis is usually faster, more cost-effective, and better applicable for laboratories that do not have immediate access to Illumina BeadChip technology.

Despite the remarkable linear correlation of age-associated DNAm with age, there is evidence for rather logarithmic association, particularly in childhood (Alisch et al. 2012; Horvath 2013; Snir et al. 2019). We selected candidate CpGs either for linear correlation (Pearson correlation), continuous non-linear association (Spearman’s rank-order), or by linear correlation with the logarithm of age. To our surprise, there was a very high overlap of selected CpGs with the three filter criteria, indicating that age-associated DNAm changes are generally accelerated in early life and are then rather linearly acquired in adulthood and the elderly. In fact, age-predictions for pediatric cohorts were clearly improved by age-transformation with a logarithmic adjustment, as elegantly described by Horvath (2013), and this approach should therefore also be considered for targeted epigenetic age predictors if applied to pediatric cohorts.

Bisulfite pyrosequencing is currently the most popular targeted method for epigenetic age predictions in forensics (Vidaki and Kayser 2018). This method is relatively simple and it has been shown to have a very high precision in DNAm measurement (Blueprint-consortium 2016). The accuracy of our epigenetic age-predictions with pyrosequencing was in a similar range as for Illumina BeadChip models. Nevertheless, the conventional PCR reaction before pyrosequencing can evoke amplification bias for methylated or non-methylated strands (Vidaki and Kayser 2018).

Droplet digital PCR might reduce this technical PCR amplification bias, because the individual droplets are either scored as positive or negative independent of the PCR efficacy. So far, only few studies used ddPCR for DNAm analysis (Pharo et al. 2018; Van Wesenbeeck et al. 2018), and only one recent study used it for measurement of age-associated DNAm changes in a pediatric cohort (Shi et al. 2018). We demonstrate that epigenetic clocks can be further enhanced with ddPCR, albeit primers and fluorescent probes could not be designed for all sequences, particularly if many neighboring CpGs were located in the target region.

Bisulfite barcoded amplicon sequencing makes use of the technical advances in massive parallel sequencing. Recently, Naue and coworkers described a similar age predictor using massive parallel sequencing (2017). Our comparative approach substantiates the notion that BBA-seq is a powerful method for epigenetic clocks, particularly if multiple samples need to be analyzed in parallel. The correlation of DNAm measurements with chronological age at individual CpGs was lower in BBA-seq data as compared to pyrosequencing or ddPCR. Furthermore, we observed a systematic off-set between the results of different sequencing runs. On the other hand, the long BBA-seq amplicons provided insight into DNAm patterns on the same DNA strand. We demonstrate that the correlation of DNAm at neighboring CpGs with chronological age follows a bell-shaped curve. Notably, this correlation often peaked at CTCF binding sites, which resembles one of the best characterized architectural proteins for the 3D chromatin conformation (Ong and Corces 2014). This is in line with a previous study indicating that age-associated DNAm changes are enriched at CTCF binding sites in old Swedish twins (Wang et al. 2018). Furthermore, it has been suggested that age-associated hypomethylation, but not hypermethylation, is associated with CTCF binding sites across various human tissues (Day et al. 2013; Reynolds et al. 2014) - albeit we observed this particularly at the hypermethylated regions in *ELOVL2*, *PDE4C* and *FHL2*. Occupancy with CTCF is tissue-specific and it is influenced by DNAm (Chang et al. 2010; Lai et al. 2010). In turn, CTCF was also capable to change DNAm status, for instance, by reducing the DNAm level at the vicinity of its binding positions (Stadler et al. 2011). Thus, the relevance of CTCF-binding for age-associated DNAm changes needs to be further explored.

Furthermore, our analysis of DNAm patterns in individual reads of BBA-seq demonstrated that neighboring CpGs are modified rather independently. We have recently described similar findings for DNAm changes during long-term culture of cells *in vitro* (Franzen et al. 2017; Franzen et al. 2018). While the stochastic changes at neighboring CpGs challenge the view of directed regulation of age-associated DNAm, they may support the notion that this process is evoked by “epigenetic drift”, possibly caused by changes in chromatin conformation. The finding of stochastic DNAm changes provided also the basis for our mathematical approach of epigenetic age predictions for individual BBA-seq reads. It is yet unclear if epigenetic aging is accelerated synchronously at different genomic regions within the same cell. Single Cell DNAm analysis has been described using WGBS (Farlik et al. 2015), but the coverage at individual CpGs (including age-associated CpGs) is notoriously very low, and thus analysis of epigenetic clocks is hampered in such datasets. On the other hand, there is evidence that age-associated DNAm is coherently modified in malignant diseases, which possibly reflects the epigenetic state of the tumor initiating cell (Lin and Wagner 2015). Thus, the epigenetic age predictions based on individual BBA-seq reads seems to reflect heterogeneity of epigenetic aging within a sample.

### Conclusions

Our comparative approach demonstrates that targeted analysis of age-associated DNAm via pyrosequencing, ddPCR, and BBA-seq enables similar precision as described for larger signatures on Illumina BeadChip profiles. The choice of regimen will rely on availability of platforms and should then be tailored for the specific application. Furthermore, our analysis of BBA-seq data demonstrated, that age-associated DNAm peaks at specific sites in the genome, particularly at CTCF binding sites. The stochastic DNAm changes at neighboring CpGs indicate, that epigenetic clocks are not directly regulated by epigenetic writers, but rather evoked by changes in higher order chromatin conformation. If we understand the underlying mechanism better, it might even be feasible to develop more precise epigenetic clocks that directly address the underlying process.

## Methods

### Sample collection

Peripheral blood samples of healthy donors (n = 131) were obtained from the Department of Transfusion Medicine at the University Hospital of RWTH Aachen. Buccal swab samples were collected with Mastaswab MD555 (Mast Group ltd., Reinfeld, Germany) at the Institute for Legal Medicine of the Heinrich Heine University in Düsseldorf, Germany (n = 95). This study has been performed according to the guidelines approved by the local ethics committees of RWTH Aachen University (EK 041/15) and Heinrich Heine University of Düsseldorf (Permit number 4939).

### Selection of age-associated CpG sites

In total 4,650 DNAm profiles of human blood samples of 12 different studies (all analyzed on the HumanMethylation450 [450k] BeadChip platform; no samples with known malignancies) were retrieved from Gene Expression Omnibus Database (GEO; Supplemental Table S1). To further select age-associated CpGs for targeted analysis we excluded i) 26,426 CpGs with high variation across the different hematopoietic subsets in (GSE35069; variances > 0.02) (Reinius et al. 2012); ii) 2,901 CpGs that are significantly affected by smoking (Gao et al. 2015; Teschendorff et al. 2015); iii) 11,648 CpGs of sex chromosomes; and iv) 31,396 CpGs that were not included in the new Illumina BeadChip EPIC platform to facilitate better comparison with future data. The remaining 416,807 CpGs were then filtered for Pearson’s correlation or Spearman correlation with chronological age. Alternatively, we analyzed the correlation with the logarithm of chronological age (Cutoffs for all comparisons: R < −0.5 or > 0.5).

Before establishing the multivariable model for epigenetic age predictions based on the 66 age-associated CpGs, we used a transformed age instead of chronological age, as described before (Horvath 2013). In brief, adult.age was assigned as 20 years, F function was used for age transformation as follows:

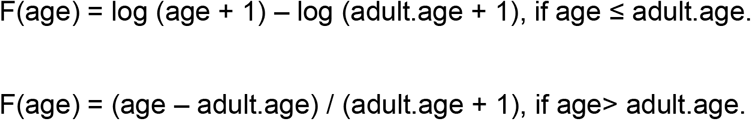

Subsequently, we established multivariable regression model (F(transformed age)) based on transformed age on the training sets from Illumina BeadChip (Supplemental Table S2). After estimation of transformed age, the results need to be inversely transformed for epigenetic age-predictions:

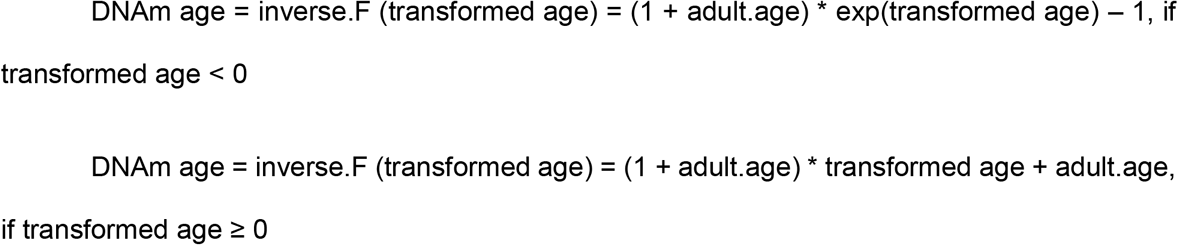

### Isolation of genomic DNA and bisulfite conversion

Genomic DNA was isolated from 50 μl blood with the QIAamp DNA Mini Kit (Qiagen, Hilden, Germany), or from buccal swab with NucleoSpin Tissue Kit (Macherey-Nagel, Düren, Germany). DNA was quantified with a Nanodrop 2000 Spectrophotometer (Thermo Scientific, Wilmington, USA) and 200 nggenomic DNA was bisulfite converted with the EZ DNA Methylation Kit (Zymo Research, Irvine, USA).

### Pyrosequencing

Bisulfite converted DNA was used for PCR amplification by PyroMark PCR Kit (Qiagen, Hilden, Germany). 20 μg of PCR products was immobilized to 5 μl Streptavidin Sepharose High Performance Bead (GE Healthcare, Piscataway, NJ, USA), and subsequently annealed to 1 μl sequencing primer (5 μM) for 2 minutes at 80°C. PCR and pyrosequencing primers (Metabion, Planegg‐Martinsried, Germany) are provided in Supplemental Figure S6 and Supplemental Table S3. Pyrosequencing was performed on PyroMark Q96 ID System and analyzed with PyroMark Q CpG software (Qiagen). To estimate epigenetic age, we either used a multivariable model based on six CpGs (Supplemental Table S4), or a nine CpG model that also considered the three CpGs of our previous work (Weidner et al. 2014) (Supplemental Table S5).

### Droplet digital PCR (ddPCR)

Droplet digital PCR was performed with a QX200™ Droplet Digital™ PCR System (Bio-Rad, CA, USA). Primers and dual-labeled probes were designed by Primer3Plus software (Supplemental Table S6). The reaction mixture consisted of 10 μl of 2X ddPCR Supermix (no dUTP; Bio-Rad), 1 μM of the forward and reverse primers, 250 nM of the probes targeting the methylated and unmethylated DNA sequences, and 25 μg of bisulfite converted DNA in a final volume of 20 μl. Together with 70 μl of droplet generation oil, it was then subjected into a DG8 disposable droplet generation cartridge (Bio-Rad). The water-in-oil droplets were produced by QX200 Droplet Generator (Bio-Rad). 40 μl of the generated droplets were transferred to the ddPCR 96-well plate (Bio-Rad). After heat sealing with the PX1 PCR Plate Sealer (Bio-Rad), the plate was placed in the C1000 Touch Thermal Cycler (Bio-Rad) for PCR run. The thermal cycling conditions were 95°C for 10 min, followed by 40 cycles of 94°C for 30 s and 1 min (2.5°C/s ramp rate) at 58°C (*FHL2* and *PDE4C)*, 54°C (*COL1A1*) or 53°C (*CCDC102B*, *MEIS1-AS3, ASPA and IGSF11*) with a 10 min step at 98 °C for enzyme deactivation and a final hold at 4 °C. Subsequently, the plates were read on the QX200 droplet reader (Bio-Rad) and data were analyzed by QuantaSoft 1.7.4 software (Bio-Rad). The percentage methylation of each reaction was calculated by Poisson statistics according to the fraction of positive droplets for methylated and non-methylated probes. The multivariable model for ddPCR is provided in Supplemental Table S7.

### Bisulfite barcoded amplicon sequencing (BBA-seq)

Target sequences with candidate CpG sites were amplified by PyroMark PCR kit (Qiagen). The forward and reverse primers contain handle sequences for the subsequent barcoding step (Supplemental Tables S8). PCR conditions were 95 °C for 15 min; 45 cycles of 94°C for 30 s, 58°C for, 72°C for 30s; and then final elongation 72°C for 10min. The amplicons of each donor (e.g. of 9 different CpGs) were pooled at equal concentrations, quantified with Qubit (Invitrogen), and cleaned up with paramagnetic beads from Agencourt AMPure PCR Purification system (Beckman Coulter). 4 μl of PCR products were subsequently added to 21 μl PyroMark Master Mix (Qiagen) containing 10 pmol of barcoded primers (adapted from NEXTflexTM 16S V1-V3 Amplicon Seq Kit, Bioo Scientific, Austin, USA) for a second PCR (95°C for 15 min; 16 cycles of 95°C for 30 s, 60°C for 30s, 72°C for 30s; final elongation 72°C for 10min). PCR products were again quantified by Qubit Kit (Invitrogen), combined in equimolar ratios, and cleaned by Select-a-Size DNA Clean & Concentrator Kit (Zymo Research, USA). 10 pM DNA library was diluted with 15% PhiX spike-in control and eventually subjected to 250 bp pair-end sequencing on a MiSeq lane (Illunima, CA, USA) using Miseq reagent V2 Nano kit (Illumina).

To estimate DNAm levels for each CpG based on BBA-seq data we used the Bismark tool (Krueger and Andrews 2011). The average number of reads per sample and genomic region was approximately 25 000 and only sequences that occurred at least 10 times were further considered. Multivariable models for epigenetic age predictions based on those CpGs that revealed highest correlation with chronological age per amplicon are provided in Supplemental Table S9 and S10 (for blood and buccal swabs, respectively). Alternative age prediction models were generated by machine learning as described before (Han et al. 2018). In brief, we applied a penalized lasso and elastic net regression model from glmnet R package on the training set from BBA-seq data. The best-fitted candidate CpGs and model was chosen by 10-fold cross validation on the training set (Supplemental Tables S11 - S14).

### Association with CTCF binding sites

Chromatin immune precipitation sequencing data (ChIP-seq) for CTCF in A549 cell lines (GSM822289), H1 human embryonic stem cells (GSM822297), and K562 cell lines (GSM822311) were analyzed. Each CpG site was extended for 250 bp in both directions and quantile normalized read counts of the ChIP-Seq experiments were compared for the 66 age-associated CpG sites in comparison to 1000 randomly chosen CpGs from the 450k array. Enrichment was estimated by Mann-Whitney rank test. The CTCF binding motifs around the target 66 age-associated CpG sites were predicted by RGT-motif analysis (www.regulatory-genomics.org/motif-analysis) with default parameters.

### Simulation of stochastic DNAm patterns

We simulated randomly generated DNAm patterns under the assumption that methylation at neighboring CpGs occurred independently. The probability that the CpG site *i* of gene *X* is methylated in the generated patterns is given by a linear function *f*_*X,S*_*(i)*, which was based on correlation of chronological age *versus* DNAm at *i* in the training set. For the simulations we used the function *random* from the *python 3* library *random*.

### Epigenetic age predictions for individual BBA-seq reads

The following algorithm was developed to estimate epigenetic age based on the binary sequel of methylated and non-methylated CpGs within individual reads of BBA-seq data: Let *X* be a gene with *n*_*X*_ CpG sites. Using the training set, we estimated the probability for each CpG to be methylated at a given age. Using linear regression, we approximated the methylation frequency of a site *i* of gene *X* as a function of age *a*. This yields a set of *n*_*X*_ linear functions *F*_*X,i*_ (*1* ≤ *i* ≤ *n*_*X*_), where *F*_*X,i*_*(a)* approximates the methylation frequency of site *i* of gene *X* at age *a*. Since the functions *F*_*X,i*_ are linear, they approach infinity or minus infinity if *a* approaches infinity or minus infinity. Since methylation frequencies assume only values between zero and one, we defined a set of *n*_*X*_ functions *p*_*X,i*_ (*1* ≤ *i* ≤ n_*X*_) as follows: *p*_*X,i*_*(a)* = *F*_*X,i*_*(a)* if *0* ≤ *F*_*X,i*_*(a)* ≤ *1*; if *F*_*X,i*_*(a)* < *0*, then *p*_*X,i*_*(a)* = *0*; if *F*_*X,i*_*(a)* > *1*, then *p*_*X,i*_*(a)* = *1*. We interpreted *p*_*X,i*_*(a)* as the probability that site *i* of gene *X* is methylated in a donor of age *a*. For a given methylation pattern *P* we could then calculate the probability *Pr(P,a)* that pattern *P* comes from a donor of age *a*. For this we assumed that methylation of different sites occurs independent from each other. Formally we obtain, *Pr(P,a)* = *q*_*1*_· … ·*q*_*nx*_, where *q*_*i*_ = *p*_*X,i*_*(a)* if site *i* is methylated in pattern *P* and *q*_*i*_ = 1 − *p*_*X,i*_*(a)* otherwise. To each detected pattern we assigned the age *a*_*P*_ that maximizes *Pr(P,a)* (if the maximum is not unique we use the average of the ages that maximize *Pr(P,a)*). The estimated age of the donor corresponded to the average of the *a*_*P*_, where we included a given value *a*_*P*_ multiple times if the pattern *P* is detected multiple times. To practically determine the *a*_*P*_, we calculated *Pr(P,a)* for *a* between 0 and 200 years in steps of one year.

## Acknowledgment

This work was supported by the German Research Foundation (DFG; WA 1706/8-1 and RI704/4-1), the Interdisciplinary Center for Clinical Research within the faculty of Medicine at the RWTH Aachen University (O3-3), Deutsche Krebshilfe (TRACK-AML), and the Federal Ministry of Education and Research (VIP + Epi-Blood-Count).

