## Supplemental figures and tables for "New Targeted Approaches for Epigenetic Age Predictions"

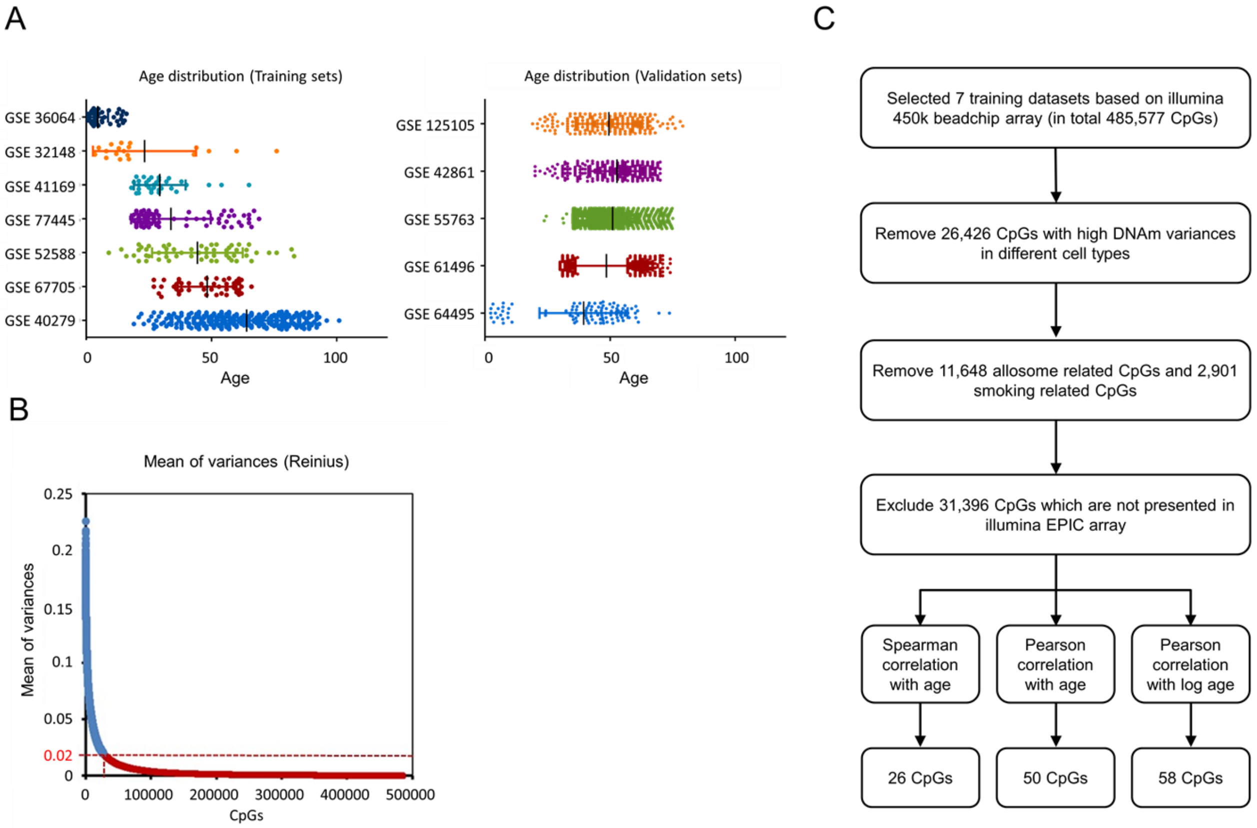

**Supplemental figure S1. Demarcation of age-associated CpGs in Illumina BeadChip datasets.**

**(A)** Age distribution of 450k Illumina BeadChip datasets in the 7 studies of the training and 5 studies of the validation cohorts (see also Supplemental Table S1; color code corresponds to Figures 1B,C, respectively). **(B)** To identify CpGs that are affected by the composition of leucocyte subsets we used the GEO datasets GSE35069 (Reinius et al. 2012). The variation of DNAm across the six leukocyte subsets was ranked and only CpGs below the threshold (R = 0.02; red dotted line) were considered for further analysis. **(C)** Schematic presentation how relevant age-associated CpGs were narrowed down. Smoking related CpGs were excluded based on two previous studies (Gao et al. 2015; Teschendorff et al. 2015).

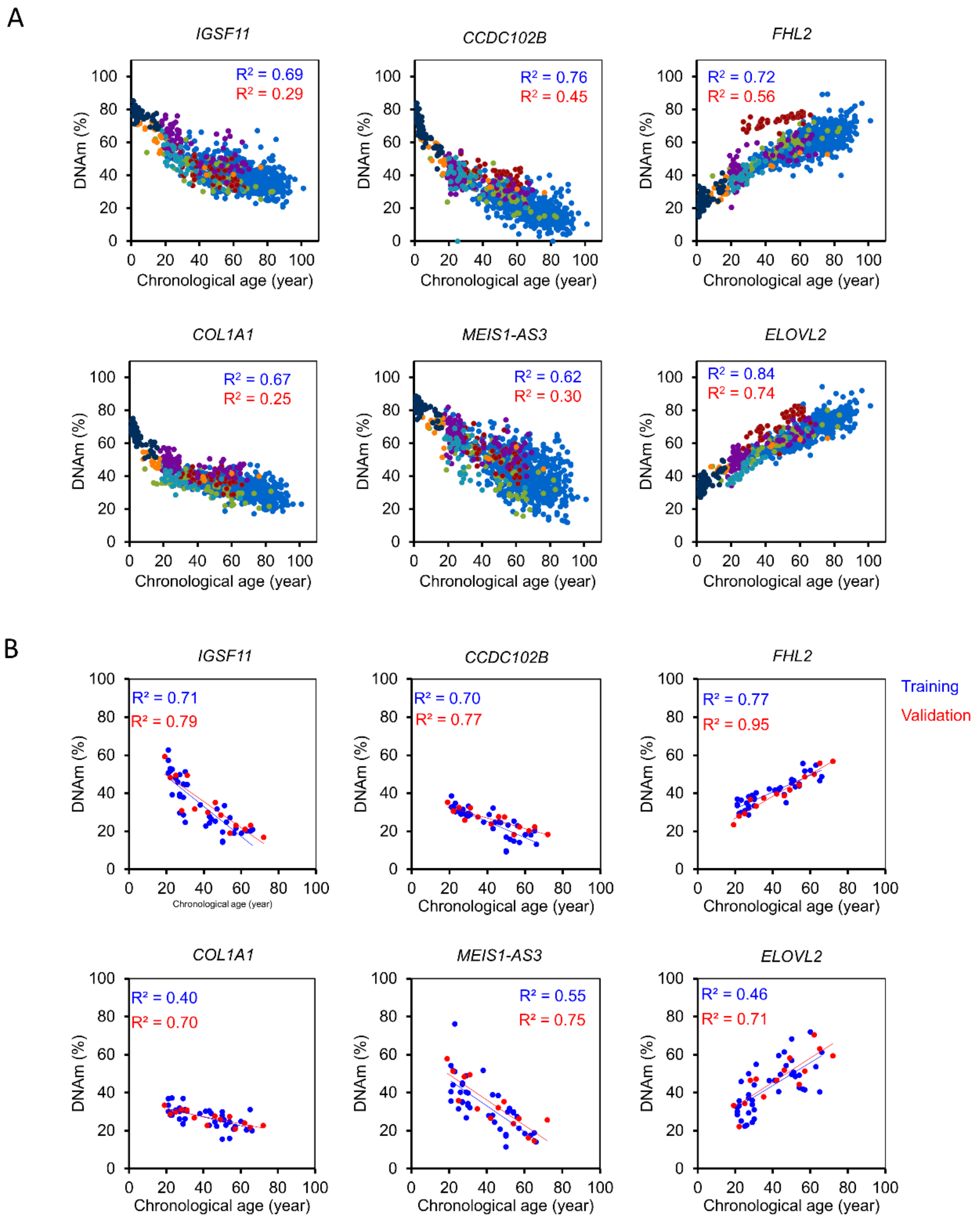

**Supplemental figure S2. Selection of 6 CpG markers for pyrosequencing.**

**(A)** DNAm levels of the Illumina BeadChip training sets were plotted for each of the 6 relevant CpGs in the genes *IGSF11*, *CCDC102B*, *FHL2*, *COL1A1*, *MEIS1-AS3* and *ELOVL2* against chronological age (color code as in Figure S1A). $R^2$ values are depicted for Illumina BeadChip training (blue) and validation sets (red). Please note that this correlation was significantly lower in the independent validation Illumina BeadChip datasets. **(B)** Pyrosequencing results for 40 blood samples of the training (blue) and 14 samples of the validation set (red).

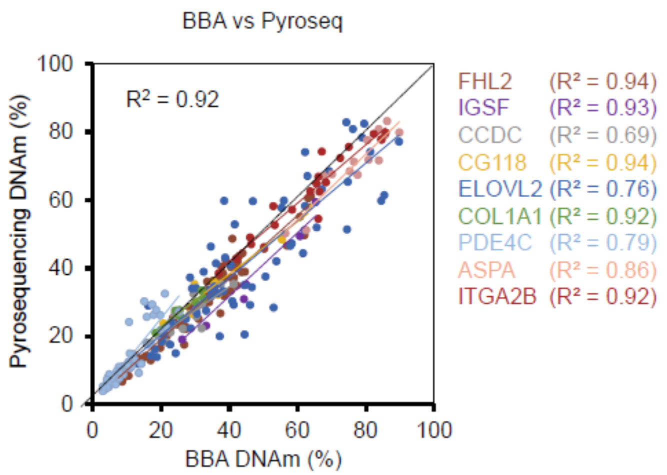

**Supplemental figure S3. Comparison of DNAm levels in pyrosequencing and BBA-seq.**

The DNA methylation levels at the nine age-associated genomic regions were analyzed in the same blood samples with bisulfite barcoded amplicon sequencing (BBA-seq) and pyrosequencing. All CpGs that were covered by both types of measurements were included into the comparison.

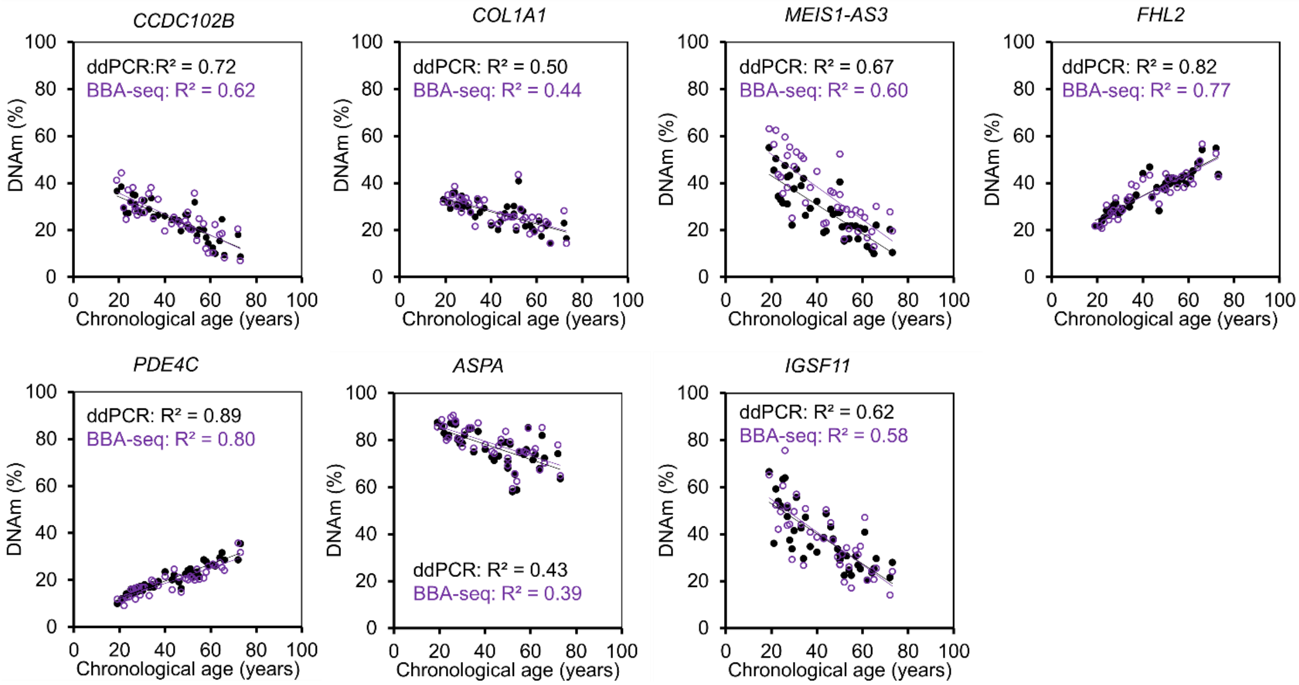

**Supplemental figure S4. Comparison of age-associated DNAm in ddPCR *versus* BBA-seq.**

DNAm measurements by droplet digital PCR (ddPCR) and bisulfite barcoded amplicon sequencing (BBA-seq) were compared for 40 blood samples. The correlation with chronological age was consistently higher in ddPCR.

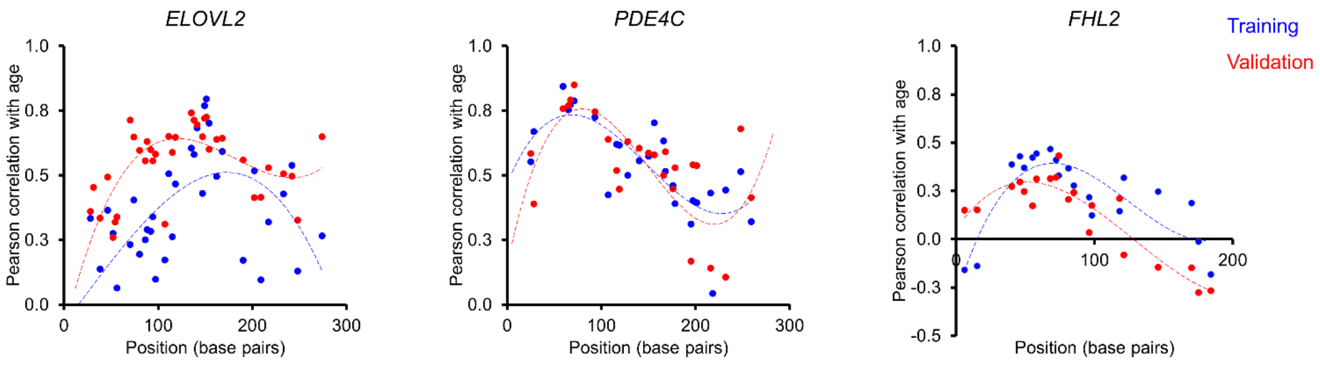

**Supplemental figure S5. Correlation of DNAm with age in buccal swabs.**

Pearson correlation of chronological age with the DNAm levels at the CpGs within amplicons of *ELOVL2*, *PDE4C* and *FHL2* in BBA-seq data of buccal swab samples.

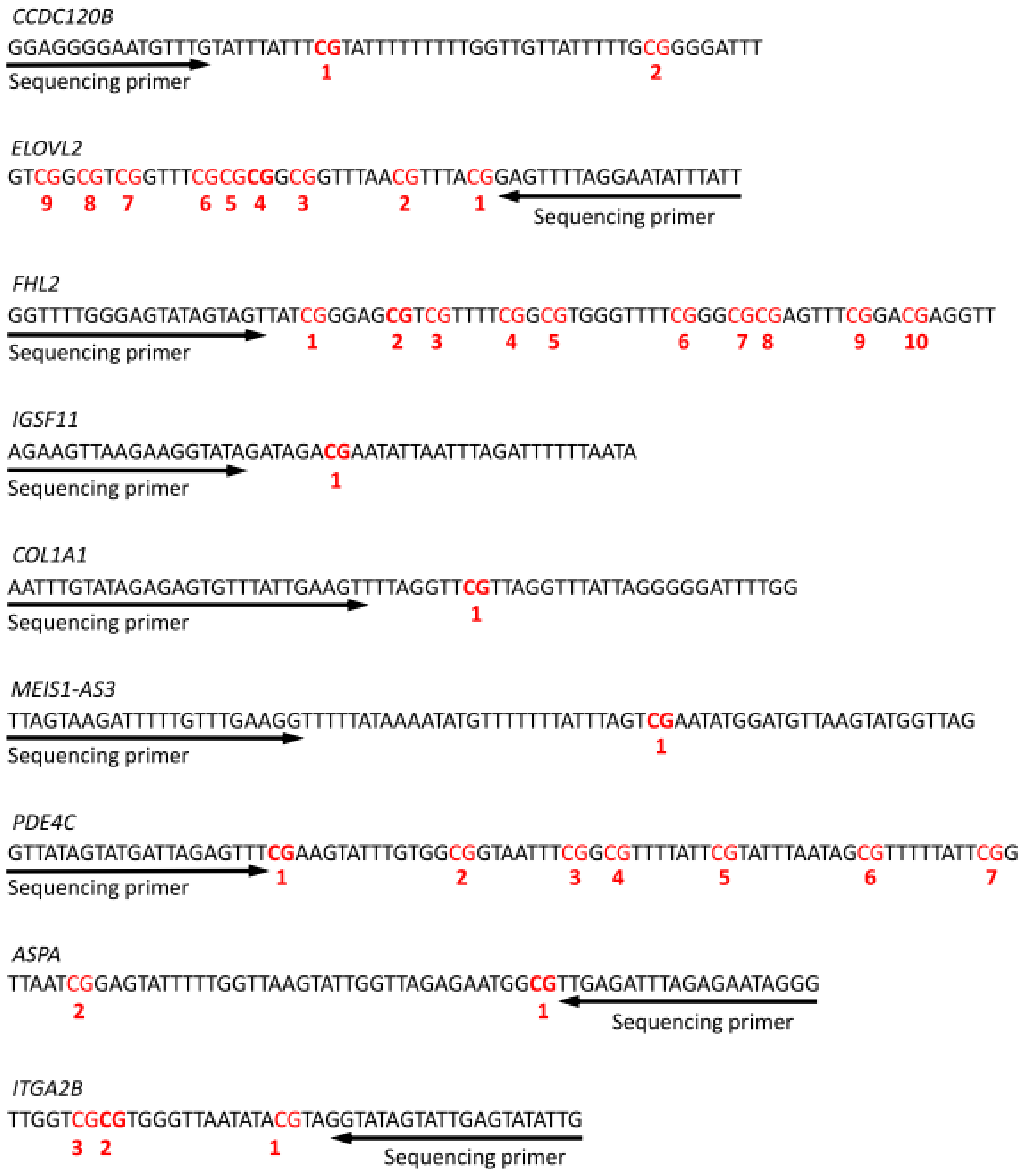

**Supplemental figure S6. Targeted sequences of pyrosequencing assays.**

Sequences for the nine genomic regions for blood aging signature are depicted and CpG sites (red) are numbered by the dispensation order of pyrosequencing. The relevant CpGs with highest age-correlation are highlighted in bold.

**1. *CCDC102B* (chr18:66,389,223-66,389,599)**

AGTGGGGCAAGTACATGATACAAGGGAGGAAACAACCGAAGTAAGGGATTGTTCTGTGAAGTAAGAGAGTATTTTGTACAGGGTTACAAGC

CTTGCCCTGATGAAGCTTTATTCAAATGGCTTCTGAGAATTTTAGATCCGTAGTATTGTCTCTGGCTTTGAACGCTGTTGAGGGAGGGGAAT
140

GTTTGCACTCATCCCGCATCCTTTTTTGGCTGCTATCTTTGCGGGGATTGTTCAAGGAGAAATCCATCCTGACTGGAATGTAGTAAAGAAAG
198

GACAGTCATTCCCAGAGAAGGGCAATTTCCCCACTCTTCCCTTGTCGTTAAATGGTCAACTGAAGCTGCAGAAATGCTGATGATACAGACA

TTACTGGCCTG

**2. *MEIS1-AS3* (chr2:66,654,583-66,654,826)**

TTTGAATAATCAGTAAGATTTTTGCCTGAAGGTTTTCACAAAATATGCCCCTTTATCCAGTCGAACATGGATGCTAAGCATGGCCAGTGTGAC

TCATCCTGTTGCTTCAAAGGTAAATGTTTGCTATTAGTGACTAGAGCACGTTCGCCAGACCTCTTCCAGCACTCAGTACTTCAACTATTAGTT

TAGAATATATCAAACGTGTATGCACTTGTGATTAGTTCCTGGAAGAATTTTCTACATT
201

**3.*COL1A1* (chr17:48,275,263-48,275,502)**

TCTGTGTGTTTGTAGAAGGAGTATGAATCTGTATAGAGAGTGCTTACTGAAGCTCCAGGCTCGCCAGGCTCACCAGGGGGACCTTGGAAG
62

CCTTGGGGACCCTTGAGAAGAAGGAAAAAGATGGGTTAGAAGACAAGTCCCTGTCAACCTTCTCCAATCTTACCAAGAGATCTCTGAGCAT

CTCTCCTGCCCTCATCCCAGTCTTCCCTCCAAAAGACCAAAGCCCGAGGAGGCATATGA

**4.Elovl2 (chr6:11,044,729-11,045,049)**

GGGGAGGGGCGCAGGGCAAGTGAGGCGGCGCCCCCCGCCCCTGCGGCCTCGCGCGCCCCCTCCTGGGCGACCGACCTCGCCCTCG
44 50 68 72

CGTCCGCGGCGTCCCCTGCCGGCCGGGCGGCGATTTGCAGGTCCAGCCGGCGCCGGTTTCGCGCGGCGGCTCAACGTCCACGGAGC
90 92 95 109 113 133 136 145147149 152

CCCAGGAATACCCACCCGCTGCCCAGATCGGCAGCCGCTGCTGCGGGGAGAAGCAGTATCGTGCAGGGCGGGCACGCTGGTCTTGCTT

ACAGTTGGGCTTCGGTGGGTTTGAAGCACACATTAGGGGGAAATGGCTCTGTTCCTGCAGGT

**5. *FHL2* (chr2:106,015,702-106,015,913)**

CTTCGTGCCCTCCGGGTCTTGGGAGCACAGTAGTTATCGGGAGCGTCGCCTCCGGCGTGGGCTCTCGGGCGCGAGTTTCGGACGAGGC
38 56

CTGGGCGCGGTGGCAGGGGTCTGCCCACGCCGGGATCTCTGCCTGGTCCCAGGAGCGGGAGACTGGAGAAGCCCCAGGACGTGCCG

GGGGAGGCGGAGGGAGGAGGGGGTCACTTCTCAGGAGG

**6.*IGSF11* (chr3:118,706,493-118,706,721)**

GCCTTGAAAAAGACAAGGAAGAAAAAATGTAGCAGCAATTCACTAGGCATGTGACTTGAGATTTGGCAGGCTGCCTTGGCAGGGCTGGAC

AGTCTGTGGGCAGAAAATTCACATATTTTTTTCCTGCAGCCAGAAGCCAAGAAGGCACAGACAGACGAATACCAACTCAGATCCTTCAACA
156

AGCTAATAATCAATAAGACCTTCCTAAGGAGAACAATGAATGAATAAC

Figure is continued on next page

**7.*ASPA* (chr17:3,379,441-3,379,591)**

ACTACTTGGTGAAATGACTTCTTGTCACATTGCTGAAGAACATATACAAAAGGTTGCTATCTTTGGAGGAACCCATGGGAATGAGCTAAC**CG**
91
GAGTATTTCTGGTTAAGCATTGGCTAGAGAATGG**CG**CTGAGATTCAGAGAACAGGGCTG
127

**8.*ITGA2B* (chr17:42,467,630-42,467,778)**

CAATCCTTTTTGGGTGATGGAGCTCTTTAACCATTAAGACTTGATTCTGGTTGGGGGCTTTGCCTAGGGGAGCCTTCCCTGACTCCTCAGG

CTGGC**CGCG**TGGGCTAACACACGTAGGCACAGCATTGAGCACACTGTTTACTCTTGGT
97 99

**9.*PDE4C* (chr19:18,343,730-18,344,002)** (reverse sequence)

CATGGAGAACCTGGGGGTCGGCGAAGGGGCAGAGGCTTGCAGCAGGTTGAGT**CG**CTCT**CGCG**GC**CG**CCACAGCATGACCAGAGCCC**C**
53 59 61 65 87
**G**AAGCACCTGTGG**CG**GCAACCCCGG**CG**CCCCATC**CG**CATCCAACAGCGCTTCTATT**CG**GATCCGGACAAGTCCGCGGGCTGCCG**CG**AG
101 113 122 144 172
AGGGACCTGAGCCCGCGGCCGGAGCTCAGGAAGTCGCGGCTCTCCTGGCCCGTTTCCTCCTGCAGGCGGTAGGTGGC**CG**GGGCAGG
253
GGCCTCCTGCAG

**Supplemental figure S7. Targeted sequences for BBA-seq.**

Sequences for nine genomic regions that were analyzed by BBA-seq are depicted and all CpG sites are highlighted in red. The age-related CpGs selected by machine learning approach are highlighted in bold and by their ordering number in the sequences (underlined CpG was used in the multivariable linear model).

### Supplemental Tables

**Table S1. DNAm profiles for candidate CpG selection**

| GEO accession ID | Set used as | Tissue | Reference | Samples | Age (years) | Gender (f/m) |
|---|---|---|---|---|---|---|
| GSE 40279 | Training | whole blood | (Hannum et al. 2013) | 656 | 19-101 | 338/318 |
| GSE 67705 | Training | Peripheral blood | (Gross et al. 2016) | 46 | 27-66 | 2/44 |
| GSE 52588 | Training | Peripheral blood | (Bacalini et al. 2015) | 58 | 9-83 | 51/7 |
| GSE 77445 | Training | Peripheral blood | (Houtepen et al. 2016) | 85 | 18-69 | 42/43 |
| GSE 41169 | Training | Peripheral blood | (Horvath and Levine 2015) | 33 | 18-65 | 12/21 |
| GSE 32148 | Training | Peripheral blood | (Harris et al. 2012) | 20 | 3.5-76 | 12/8 |
| GSE 36064 | Training | Peripheral blood | (Alisch et al. 2012) | 78 | 1-15.3 | 0/78 |
| GSE 64495 | Validation | Peripheral blood | (Walker et al. 2015) | 106 | 2.3-73.7 | 76/37 |
| GSE 61496 | Validation | Peripheral blood | (Tan et al. 2014) | 312 | 30-74 | 148/164 |
| GSE 55763 | Validation | Peripheral blood | (Lehne et al. 2015) | 2711 | 23.7-75 | 859/1805 |
| GSE 42861 | Validation | Peripheral blood | (Liu et al. 2013) | 335 | 20-70 | 239/96 |
| GSE125105 | Validation | Peripheral blood | - | 210 | 19-79 | 126/84 |

All datasets were generated on the 450k Illumina BeadChip. Data are accessible under http://www.ncbi.nlm.nih.gov/geo/.

**Table S2. Multivariable 66 CpG model for Illumina BeadChip data**

| Target site | Gene Name | CHR | Map Info | Coeffients |
|---|---|---|---|---|
| (Intercept) | | | | 0.70 |
| cg19283806 | *CCDC102B* | 18 | 66389420 | -0.59 |
| cg11807280 | *MEIS1-AS3* | 2 | 66654644 | -0.21 |
| cg00329615 | *IGSF11* | 3 | 118706648 | 0.01 |
| cg22454769 | *FHL2* | 2 | 106015767 | 0.05 |
| cg16867657 | *ELOVL2* | 6 | 11044877 | 2.16 |
| cg22796704 | *ARHGAP22* | 10 | 49673534 | -0.69 |
| cg09809672 | *EDARADD* | 1 | 236557682 | -0.65 |
| cg18618815 | *COL1A1* | 17 | 48275324 | -0.76 |
| cg25533247 | *AKAP8L* | 19 | 15530630 | 0.13 |
| cg02286081 | *HLA-DPB1* | 6 | 33043841 | -0.64 |
| cg20222376 | *AKAP8L* | 19 | 15530606 | 0.02 |
| cg19344626 | *NWD1* | 19 | 16830749 | -0.06 |
| cg07082267 | | 16 | 85429035 | -0.23 |
| cg26350754 | *HLA-DPB1* | 6 | 33043868 | 0.51 |
| cg15845821 | *NWD1* | 19 | 16830613 | 1.57 |
| cg11741201 | *FJX1* | 11 | 35638398 | 0.35 |
| cg16054275 | *F5* | 1 | 169556022 | 0.19 |
| cg18933331 | | 1 | 110186418 | -0.42 |
| cg20249566 | *NWD1* | 19 | 16830739 | -0.78 |
| cg16604658 | *TBK1* | 12 | 64847188 | 0.48 |
| cg07583137 | *CHMP4C* | 8 | 82644012 | 0.05 |
| cg16008966 | | 1 | 114761794 | -0.39 |
| cg14556683 | *EPHX3* | 19 | 15342982 | 0.10 |
| cg03746976 | *C16orf57* | 16 | 58035805 | 0.02 |
| cg14314729 | | 5 | 179815975 | 0.29 |
| cg03431918 | | 17 | 77716367 | -0.27 |
| cg22156456 | *EIF1* | 17 | 39844239 | 0.26 |
| cg23078123 | *GPR177* | 1 | 68577796 | -0.74 |
| cg09748749 | *ASL* | 7 | 65540429 | -1.16 |
| cg17457912 | *C17orf91* | 17 | 1617102 | -0.03 |
| cg06492796 | | 12 | 96883057 | -0.18 |
| cg17593342 | | 6 | 14037614 | 0.85 |
| cg05308819 | | 1 | 155959156 | -0.87 |
| cg22512670 | *RPS6KA1* | 1 | 26855765 | -0.32 |
| cg01820962 | *NT5DC1* | 6 | 116511817 | -0.85 |
| cg06639320 | *FHL2* | 2 | 106015739 | 1.94 |
| cg03224418 | *SAMD10;PRPF6* | 20 | 62611858 | 0.91 |
| cg17436656 | *RARG* | 12 | 53627106 | -0.10 |

| | | | | |
|---|---|---|---|---|
| cg19500607 | *HTR4* | 5 | 148034319 | 0.52 |
| cg03735592 | *NHSL1* | 6 | 138821354 | 0.33 |
| cg20669012 | | 3 | 11102341 | 0.36 |
| cg19761273 | *CSNK1D* | 17 | 80232096 | 0.01 |
| cg07080372 | *SLC25A22* | 11 | 796607 | -1.46 |
| cg03638795 | *SIGIRR* | 11 | 416499 | -0.28 |
| cg19722847 | *IPO8* | 12 | 30849114 | -0.59 |
| cg24711336 | | 10 | 80063791 | 0.65 |
| cg26935102 | *POLR3GL;ANKRD34A* | 1 | 145470946 | 0.50 |
| cg10221746 | | 1 | 156629412 | 0.24 |
| cg02085953 | *ARID5A* | 2 | 97202260 | -0.63 |
| cg04604946 | *LRRC23* | 12 | 7023352 | -1.10 |
| cg08558886 | | 2 | 151469837 | -0.15 |
| cg22361181 | *NKIRAS2* | 17 | 40171740 | 0.62 |
| cg04208403 | *ZNF423* | 16 | 49525807 | -0.09 |
| cg12623930 | *ABHD14B* | 3 | 52008802 | 0.04 |
| cg21572722 | *ELOVL2* | 6 | 11044894 | -0.49 |
| cg17885226 | | 6 | 105388731 | 0.35 |
| cg00748589 | | 12 | 11653486 | 2.53 |
| cg13033938 | *IP6K1* | 3 | 49824475 | -3.64 |
| cg19784428 | *NWD1* | 19 | 16830746 | 0.08 |
| cg22016779 | *DNER* | 2 | 230452311 | -0.44 |
| cg01974375 | *PI4KB* | 1 | 151298954 | -0.40 |
| cg25256723 | *F5* | 1 | 169555944 | -0.26 |
| cg24724428 | *ELOVL2* | 6 | 11044888 | 0.72 |
| cg07547549 | *SLC12A5* | 20 | 44658225 | -0.10 |
| cg25410668 | *RPA2* | 1 | 28241577 | 0.28 |
| cg21296230 | *GREM1* | 15 | 33010536 | 0.93 |

This model was trained for transformed age instead of chronological age, as described before (Horvath 2013).

**Table S3. Primer list for Pyrosequencing assays**

| Primer | Sequence |
|---|---|
| ***CCDC102B*** | |
| Forward | 5'- TGTTGAGGGAGGGGAATGTTTGTATTTAT-3' |
| Reverse | 5'-Biotin- CCAATAATATCTATATCATCAACATTTCTACAACTT-3' |
| Sequencing | 5'- GGAGGGGAATGTTTG -3' |
| ***IGSF11*** | |
| Forward | 5'- GTTGGATAGTTTGTGGGTAGAAAATTTA -3' |
| Reverse | 5'-Biotin- ATTATTCATTCATTATTCTCCTTAAAAAAATCTTATT -3' |
| Sequencing | 5'- AGAAGTTAAGAAGGTATAGATA -3' |
| ***ELOVL2*** | |
| Forward | 5'-Biotin- GGGAGGGGAGTAGGGTAAGTGA -3' |
| Reverse | 5'- CCATCTAAACAACCAATAAATATTCCTAAAAC -3' |
| Sequencing | 5'- AATAAATATTCCTAAAACTC -3' |
| ***COL1A1*** | |
| Forward | 5'- TTGAAGGGAAGAGGTAAGGAAGATTTTA -3' |
| Reverse | 5'- Biotin- TAACCCATCTTTTTCCTTCTTCTCA -3' |
| Sequencing | 5'- AATTTGTATAGAGAGTGTTTATTG -3' |
| ***MEIS1-AS3*** | |
| Forward | 5'- TTGAATAATTAGTAAGATTTTTGTTTGAAGGTTT -3' |
| Reverse | 5'-Biotin- TTACCTTTAAAACAACAAAATAAATCACACTAACC -3' |
| Sequencing | 5'- TTAGTAAGATTTTTGTTTG -3' |
| ***FHL2*** | |
| Forward | 5'- GTGTTTTTAGGGTTTTGGGAGTATAGTAGT -3' |
| Reverse | 5'-Biotin- CACCTCCTAAAACTTCTCCAATCTCC -3' |
| Sequencing | 5'-TATTTTTTAAGGTAGTAAGAGT-3' |
| ***ASPA*** | |
| Forward | 5'-Biotin- ATTATTTGGTGAAATGATT -3' |
| Reverse | 5'- CAACCCTATTCTCTAAATCTC -3' |
| Sequencing | 5'- CCCTATTCTCTAAATCTCA -3' |
| ***ITGA2B*** | |
| Forward | 5'-Biotin- TAATTTTTTTTGGGTGATG -3' |
| Reverse | 5'- ACCAAAAATAAACAATATACTCAAT -3' |
| Sequencing | 5'- CAATATACTCAATACTATACCT -3' |
| ***PDE4C*** | |
| Forward | 5'- AGGTTTGTAGTAGGTTGAG -3' |
| Reverse | 5'-Biotin- AACTCAAATCCCTCTC -3' |
| Sequencing | 5'- GTTATAGTATGATTAGAGTTT -3' |

**Table S4. Epigenetic aging signature based on pyrosequencing (6 CpG model)**

| Target site | Gene Name | CHR | Map Info | Coeffients |
|---|---|---|---|---|
| (Intercept) | | | | 24.78 |
| cg22454769 | *FHL2* | 2 | 106015767 | 0.86 |
| cg00329615 | *IGSF11* | 3 | 118706648 | -0.22 |
| cg19283806 | *CCDC102B* | 18 | 66389420 | -0.26 |
| cg11807280 | *MEIS1-AS3*[1] | 2 | 66654644 | -0.17 |
| cg16867657 | *ELOVL2* | 6 | 11044877 | 0.10 |
| cg18618815 | *COL1A1* | 17 | 48275324 | -0.17 |

**Table S5. Epigenetic aging signature based on pyrosequencing (9 CpG model)**

| Target site | Gene Name | CHR | Map Info | Coeffients |
|---|---|---|---|---|
| (Intercept) | | | | 5.00 |
| cg22454769 | *FHL2* | 2 | 106015767 | 0.63 |
| cg00329615 | *IGSF11* | 3 | 118706648 | -0.17 |
| cg19283806 | *CCDC102B* | 18 | 66389420 | -0.05 |
| cg11807280 | *MEIS1-AS3*[1] | 2 | 66654644 | -0.11 |
| cg16867657 | *ELOVL2* | 6 | 11044877 | 0.15 |
| cg18618815 | *COL1A1* | 17 | 48275324 | -0.04 |
| cg02228185 | *ASPA* | 17 | 3379567 | -0.05 |
| cg25809905 | *ITGAB* | 17 | 42467728 | 0.04 |
| NA[1] | *PDE4C* | 19 | 18343915 | 0.74 |

[1] A neighboring CpG was used, which is not included on the Illumina BeadChip.

**Table S6. Primer list for ddPCR assay**

| Primer | Sequence |
|---|---|
| ***CCDC102B*** | |
| Forward | 5'- AGTATTGTTTTTGGTTTTGAA-3' |
| Reverse | 5'- CCCTTCTCTAAAAATAACTATCC-3' |
| Probe | 6-Fam - AGGGAGGGGAATGTTTGTATTTATTTCGTA -BHQ-1[a] |
| | Hex-AGGGAGGGGAATGTTTGTATTTATTTTGTA -BHQ-1[b] |
| ***COL1A1*** | |
| Forward | 5'-AGGAGAGTTTGTGTGTTTGT-3' |
| Reverse | 5'-TCTAACCCATCTTTTTCCTTCT-3' |
| Probe | 6-Fam -TGAAGTTTTAGGTTCGTTAGGTTTATTAGG - BHQ-1[a] |
| | Hex -TGAAGTTTTAGGTTTGTTAGGTTTATTAGG - BHQ-1[b] |
| ***MEIS1-AS3*** | |
| Forward | 5'-AGAGTAYGTTYGTTAGATTT-3' |
| Reverse | 5'- AAATCCTCATAACAATAACTTAAAA-3' |
| Probe | 6-Fam - AGAATATATTAAACGTGTATGTATTTGTGA - BHQ-1[a] |
| | Hex - AGAATATATTAAATGTGTATGTATTTGTGA - BHQ-1[b] |
| ***FHL 2*** | |
| Forward | 5'-TATTTTTTGTTTGTTAGGGTTTTT-3' |
| Reverse | 5'-TCCTAAAACCAAACAAAAATCC-3' |
| Probe | 6-Fam -TTTTGGGAGTATAGTAGTTATCGGGAG - BHQ-1[a] |
| | Hex -TTTTGGGAGTATAGTAGTTATTGGGAG - BHQ-1[b] |
| ***ASPA*** | |
| Forward | 5'-AGGTTGTTATTTTTGGAGGA-3' |
| Reverse | 5'-CCTCCAACCCTATTCTCTAA-3' |
| Probe | 6-Fam -TGGGAATGAGTTAATCGGAGTAT- BHQ-1[a] |
| | Hex -TGGGAATGAGTTAATTGGAGTAT- BHQ-1[b] |
| ***IGSF11*** | |
| Forward | 5'-TGGTAGGGTTGGATAGTT-3' |
| Reverse | 5'-AATTATTCATTCATTATTCTCCTTAA-3' |
| Probe | 6-Fam -AAGGTATAGATAGACGAATATTAATTTAGA- BHQ-1[a] |
| | Hex -AAGGTATAGATAGATGAATATTAATTTAGA- BHQ-1[b] |
| ***PDE4C*** | |
| Forward | 5'-GAGGTTTGTAGTAGGTTGAGT-3' |
| Reverse | 5'-CRAACTCAAATCCCTCTCR-3' |
| Probe | 6-Fam -TAGTATGATTAGAGTTTCGAAGTATTTGTG- BHQ-1[a] |
| | Hex -TAGTATGATTAGAGTTTTGAAGTATTTGTG- BHQ-1[b] |

[a] Probe targeting the methylated sequence.
[b] Probe targeting the non-methylated sequence.

**Table S7. Multivariable model for ddPCR (7 CpG)**

| Target site | Gene Name | CHR | Map Info | Coeffients |
|---|---|---|---|---|
| (Intercept) | | | | 16.49 |
| cg06639320 | *FHL2* | 2 | 106015739 | 1.28 |
| cg00329615 | *IGSF11* | 3 | 118706648 | 0.11 |
| cg19283806 | *CCDC102B* | 18 | 66389420 | 0.03 |
| NA[1] | *MEIS1-AS3*[1] | 2 | 66654782 | 0.02 |
| cg18618815 | *COL1A1* | 17 | 48275324 | -0.59 |
| NA[1] | *ASPA* | 17 | 3379531 | -0.28 |
| NA[1] | *PDE4C* | 19 | 18343915 | 0.57 |

[1] A neighboring CpG was used, which is not included on the Illumina BeadChip.

**Table S8. Primer list for BBA-seq assay**

| Primer | Sequence |
|---|---|
| ***CCDC102B*** | |
| Forward | 5'-CTCTTTCCCTACACGACGCTCTTCCGATCTAGTGGGGTAAGTATATGATATAAGGGAGGAAATA -3' |
| Reverse | 5'- CTGGAGTTCAGACGTGTGCTCTTCCGATCTCAAACCAATAATATCTATATCATCAACATTTCT -3' |
| ***IGSF11*** | |
| Forward | 5'- CTCTTTCCCTACACGACGCTCTTCCGATCTGTTTTGAAAAAGATAAGGAAGAAAAAATGTAGTA-3' |
| Reverse | 5'- CTGGAGTTCAGACGTGTGCTCTTCCGATCTATTATTCATTCATTATTCTCCTTAAAAAAATCTTATT -3' |
| ***ELOVL2*** | |
| Forward | 5'- CTCTTTCCCTACACGACGCTCTTCCGATCTGGGGAGGGGAGTAGGGTAAGTGA-3' |
| Reverse | 5'- CTGGAGTTCAGACGTGTGCTCTTCCGATCTACCTACAAAAACAAAACCATTTCCCCCTAATAT-3' |
| ***COL1A1*** | |
| Forward | 5'- CTCTTTCCCTACACGACGCTCTTCCGATCTTTTGTGTGTTTGTAGAAGGAGTATGAATTTGTATAG -3' |
| Reverse | 5'- CTGGAGTTCAGACGTGTGCTCTTCCGATCTTCATATACCTCCTCCAACTTTAATCTTTTAAA-3' |
| ***MEIS1-AS3*** | |
| Forward | 5'- CTCTTTCCCTACACGACGCTCTTCCGATCTTTTGAATAATTAGTAAGATTTTTGTTTGAAGGTTT-3' |
| Reverse | 5'- CTGGAGTTCAGACGTGTGCTCTTCCGATCTAATATAAAAAATTCTTCCAAAAACTAATCACAAATACA -3' |
| ***FHL2*** | |
| Forward | 5'- CTCTTTCCCTACACGACGCTCTTCCGATCTTTTAGTGTTTTTAGGGTTTTGGGAGTATAGTAGTT-3' |
| Reverse | 5'- CTGGAGTTCAGACGTGTGCTCTTCCGATCTCCTCCTAAAAAATAACCCCCTCCTCCCT-3' |
| ***ASPA*** | |
| Forward | 5'-CTCTTTCCCTACACGACGCTCTTCCGATCTATTATTTGGTGAAATGATT-3' |
| Reverse | 5'- CTGGAGTTCAGACGTGTGCTCTTCCGATCTCAACCCTATTCTCTAAATCTC -3' |
| ***ITGA2B*** | |
| Forward | 5'-CTCTTTCCCTACACGACGCTCTTCCGATCTTAATTTTTTTTGGGTGATG -3' |
| Reverse | 5'- CTGGAGTTCAGACGTGTGCTCTTCCGATCTACCAAAAATAAACAATATACTCAAT-3' |
| ***PDE4C*** | |
| Forward | 5'- CTCTTTCCCTACACGACGCTCTTCCGATCTTATGGAGAATTTGGGG -3' |
| Reverse | 5'- CTGGAGTTCAGACGTGTGCTCTTCCGATCTCTACAAAAAACCCCTACC -3' |
| ***CD6*** | |
| Forward | 5'- CTCTTTCCCTACACGACGCTCTTCCGATCTAGTATAGGTAGTTGGGGTTTTTTTTATTAGTTTTTGTA-3' |
| Reverse | 5'- CTGGAGTTCAGACGTGTGCTCTTCCGATCTCCAAATCTACTCTACCCTTTACTATTCTTATTCCTAT-3' |
| ***SERPINB5*** | |
| Forward | 5'- CTCTTTCCCTACACGACGCTCTTCCGATCTATTGTGGATAAGTTGTTAAGAGGTTTGAGTAGG-3' |
| Reverse | 5'- CTGGAGTTCAGACGTGTGCTCTTCCGATCTAAACAAACAAACCAAAAACACAAAAACCTAAATAT-3' |

**Table S9. Multivariable model for BBA-seq of blood (9 CpG model)**

| Target site | Gene Name | CHR | Map Info | Coeffients |
|---|---|---|---|---|
| (Intercept) | | | | 22.57 |
| NA[1] | *FHL2* | 2 | 106015747 | 0.68 |
| cg00329615 | *IGSF11* | 3 | 118706648 | -0.09 |
| cg19283806 | *CCDC102B* | 18 | 66389420 | 0.07 |
| NA[1] | *MEIS1-AS3* | 2 | 66654782 | 0.26 |
| cg16867657 | *ELOVL2* | 6 | 11044877 | 0.53 |
| cg18618815 | *COL1A1* | 17 | 48275324 | -0.01 |
| NA[1] | *ASPA* | 17 | 3379531 | 0.69 |
| NA[1] | *ITGAB* | 17 | 42467726 | -0.39 |
| NA[1] | *PDE4C* | 19 | 18343886 | -0.41 |

[1] A neighboring CpG was used, which is not included on the Illumina BeadChip.

**Table S10. Multivariable model for BBA-seq of buccal swabs (9 CpG model)**

| Target site | Gene Name | CHR | Map Info | Coeffients |
|---|---|---|---|---|
| (Intercept) | | | | 24.10 |
| NA[1] | *FHL2* | 2 | 106015747 | 0.68 |
| cg00329615 | *IGSF11* | 3 | 118706648 | -0.07 |
| cg19283806 | *CCDC102B* | 18 | 66389420 | 0.07 |
| NA[1] | *MEIS1-AS3* | 2 | 66654782 | 0.26 |
| cg16867657 | *ELOVL2* | 6 | 11044877 | 0.53 |
| cg18618815 | *COL1A1* | 17 | 48275324 | -0.02 |
| NA[1] | *ASPA* | 17 | 3379531 | 0.69 |
| NA[1] | *ITGAB* | 17 | 42467726 | -0.40 |
| NA[1] | *PDE4C* | 19 | 18343886 | -0.42 |

[1] A neighboring CpG was used, which is not included on the Illumina BeadChip.

**Table S11. Machine learning model by Lasso (blood)**

| | Target sites* | Coefficients |
|---|---|---|
| | (Intercept) | 14.02 |
| 1 | ASPA.093 | -0.15 |
| 2 | CCDC102B.142 | -0.38 |
| 3 | CCDC102B.200 | -0.04 |
| 4 | CG11807280.203 | -0.09 |
| 5 | COL1A1.063 | -0.04 |
| 6 | Elovl2.074 | 0.10 |
| 7 | Elovl2.092 | 0.08 |
| 8 | Elovl2.115 | 0.28 |
| 9 | Elovl2.151 | 0.44 |
| 10 | FHL2.146 | 0.39 |
| 11 | ITGA2B.099 | -0.11 |
| 12 | PDE4C.059 | 0.16 |
| 13 | PDE4C.067 | 0.24 |
| 14 | PDE4C.071 | 0.18 |
| 15 | PDE4C.093 | 0.30 |
| 16 | PDE4C.119 | 0.33 |
| 17 | PDE4C.128 | 0.13 |

* Target sites correspond to the CpG# in Figure S7.

**Table S12. Machine learning model by ElasticNet (blood)**

| | Target sites* | Coefficients |
|---|---|---|
| | (Intercept) | 17.46 |
| 1 | ASPA.093 | -0.16 |
| 2 | CCDC102B.142 | -0.35 |
| 3 | CCDC102B.200 | -0.01 |
| 4 | CG11807280.064 | -0.02 |
| 5 | CG11807280.203 | -0.07 |
| 6 | COL1A1.063 | -0.07 |
| 7 | Elovl2.070 | 0.39 |
| 8 | Elovl2.092 | 0.03 |
| 9 | Elovl2.115 | 0.26 |
| 10 | Elovl2.135 | 0.08 |
| 11 | Elovl2.138 | 0.03 |
| 12 | Elovl2.147 | 0.02 |
| 13 | Elovl2.149 | 0.04 |
| 14 | Elovl2.151 | 0.25 |
| 15 | Elovl2.154 | 0.06 |
| 16 | FHL2.040 | 0.02 |
| 17 | FHL2.058 | 0.02 |
| 18 | FHL2.146 | 0.34 |
| 19 | ITGA2B.099 | -0.15 |
| 20 | PDE4C.059 | 0.14 |
| 21 | PDE4C.065 | 0.10 |
| 22 | PDE4C.067 | 0.20 |
| 23 | PDE4C.071 | 0.17 |
| 24 | PDE4C.093 | 0.25 |
| 25 | PDE4C.119 | 0.32 |
| 26 | PDE4C.128 | 0.14 |

* Target sites correspond to the CpG# in Figure S7.

**Table S13. Machine learning model by Lasso (swab)**

| | Target sites* | Coefficients |
|---|---|---|
| | (Intercept) | -15.27 |
| 1 | ASPA.129 | -0.16 |
| 2 | CCDC102B.323 | -0.06 |
| 3 | CG11807280.144 | 0.17 |
| 4 | CG11807280.203 | 0.08 |
| 5 | Elovl2.046 | 0.06 |
| 6 | Elovl2.052 | 0.64 |
| 7 | Elovl2.094 | -0.06 |
| 8 | Elovl2.097 | -0.07 |
| 9 | Elovl2.111 | 0.03 |
| 10 | Elovl2.135 | 0.32 |
| 11 | Elovl2.149 | 0.29 |
| 12 | Elovl2.151 | 0.12 |
| 13 | Elovl2.154 | 0.10 |
| 14 | Elovl2.202 | 0.02 |
| 15 | Elovl2.233 | 0.05 |
| 16 | Elovl2.248 | -0.01 |
| 17 | FHL2.058 | 0.11 |
| 18 | FHL2.072 | 0.42 |
| 19 | ITGA2B.099 | -0.12 |
| 20 | ITGA2B.101 | -0.03 |
| 21 | PDE4C.059 | 0.12 |
| 22 | PDE4C.065 | 0.30 |
| 23 | PDE4C.107 | 0.22 |
| 24 | PDE4C.119 | 0.64 |
| 25 | PDE4C.150 | 0.21 |
| 26 | PDE4C.178 | 0.39 |
| 27 | PDE4C.259 | 0.29 |

* Target sites correspond to the CpG# in Figure S7.

**Table S14. Machine learning model by ElasticNet (swab)**

| | Target sites* | Coefficients |
|---|---|---|
| | (Intercept) | -10.35 |
| 1 | ASPA.129 | -0.15 |
| 2 | CCDC102B.323 | -0.06 |
| 3 | CG11807280.144 | 0.16 |
| 4 | CG11807280.203 | 0.01 |
| 5 | Elovl2.052 | 0.63 |
| 6 | Elovl2.111 | 0.04 |
| 7 | Elovl2.135 | 0.27 |
| 8 | Elovl2.149 | 0.26 |
| 9 | Elovl2.151 | 0.16 |
| 10 | Elovl2.154 | 0.10 |
| 11 | Elovl2.202 | 0.0001 |
| 12 | Elovl2.233 | 0.03 |
| 13 | FHL2.058 | 0.17 |
| 14 | FHL2.072 | 0.28 |
| 15 | ITGA2B.099 | -0.13 |
| 16 | ITGA2B.101 | -0.03 |
| 17 | PDE4C.059 | 0.15 |
| 18 | PDE4C.065 | 0.27 |
| 19 | PDE4C.067 | 0.02 |
| 20 | PDE4C.071 | 0.004 |
| 21 | PDE4C.093 | 0.02 |
| 22 | PDE4C.107 | 0.25 |
| 23 | PDE4C.119 | 0.57 |
| 24 | PDE4C.150 | 0.19 |
| 25 | PDE4C.178 | 0.34 |
| 26 | PDE4C.259 | 0.19 |

* Target sites correspond to the CpG# in Figure S7.
